# A critical role of protein damage response in mediating cancer drug resistance

**DOI:** 10.1101/2024.07.17.603840

**Authors:** Fangyuan Shao, Jie Li, Hao Xiao, Ling Li, Bo Li, YuJun Chen, Xiangpeng Chu, Maoxin Ran, Dongyang Tang, Yuzhong Peng, Yujian Huang, Lijian Wang, Yanxia Shi, Nan Shao, Kai Miao, Changhua Zhang, Ying Lin, Jun Yan, Kin Yip Tam, Xiaoling Xu, Chu-Xia Deng

## Abstract

Multidrug resistance (MDR) frequently occurs during cancer therapy and remains a major obstacle for the cure of most cancers(*1, 2*). Drug resistance could exist intrinsically or be acquired by drug treatment(*3–5*), yet factors that regulate the resistance remain elusive. Here, we show that most anticancer drugs damage neosynthesized proteins prior to reaching their canonic targets and elicit profound cytotoxicity, which is largely compensated by protein damage response (PDR). We demonstrate that the PDR includes damage recognition and clearance that are mainly mediated by ubiquitin and proteasome systems, although some other factors, including cellular ATP levels and proliferation status, are also involved. We show that cancer stem cells (CSCs), which have lower protein synthesis, and drug resistance acquired cells (DRAC), which have higher proteasome activity, are more resistant than other cells. We further demonstrate that ATP promotes protein synthesis and suppresses proteasome activity, thus, increasing mitochondrial ATP production by PDK1 inhibition and using proteasome inhibitor to block protein damage clearance render CSCs and DRACs more vulnerable to anticancer drugs. Thus, patients with drug-resistant cancers and treatment-naïve patients with low ATP levels and/or high proteasome activity can be identified and subtyped, and therapies containing PDK1-I and/or proteasome-I may be effective options for these patients.

## Introduction

According to estimates by the International Agency for Research on Cancer, there were 19.3 million new cancer cases and 9.96 million cancer deaths worldwide in 2020. One of the main reasons for the high cancer mortality rate is that many patients develop resistance to drugs during drug treatment, which leads to drug treatment failure or reduced curative effects and aggravates the development of tumors (*6–8*).

Some patients exhibit congenital drug resistance; that is, these patients have no response to anticancer drugs at the beginning of treatment. In many other cases, drug resistance is acquired during the treatment process, i.e., patients show a good therapeutic effect in the beginning but gradually lose responsiveness during chemotherapy (*6*). Even more difficult to treat, after clinical patients acquire tolerance to therapeutic drugs, they often become cross-resistant to other drugs that they have not been exposed to with different structures and different mechanisms of action, a phenomenon called multidrug resistance (MDR) (*7*).

Recent studies have shown that the mechanisms of MDR (*8*) involve DNA damage repair, drug target mutation, drug metabolism and inactivation, cellular drug excretion, aberrant cell death regulation, cancer stem cells (CSCs), etc. (*7*). To identify specific targets and pathways that may overcome drug resistance, we previously utilized RNAi-mediated gene knockdown and high-throughput drug library screening in multiple cancer cell lines to identify top candidates that were then subjected to in vivo validation (*3, 9, 10*). Using a whole-genome RNAi library containing >22,000 genes, we found that cisplatin damages both DNA and proteins and that drug resistance is closely related to the ability of cancer cells to remove damaged proteins, which is accompanied by increased proteasome activity and reduced mitochondrial respiratory activity (*3*). The same data were obtained in models of cancer drug resistance evolution generated by an increasing cisplatin concentration gradient (*3*). We also found that once the cells became cisplatin resistant, they were also cross-resistant to > 40 anticancer drugs; notably, this MDR could be reversed by blocking proteasome activity to block the clearance of damaged proteins, suggesting that protein damage could serve as a common toxic mechanism by which anticancer drugs kill cancer cells. While this study links anticancer drugs with protein damage and MDR with an enhanced ability to remove damaged proteins, it leaves several questions unanswered, including how often and under what conditions anticancer drugs damage proteins, the mechanism by which resistant cancer cells evade drug-induced protein damage, and how this mechanism can be overcome.

In this study, we find that most anticancer drugs tested can cause protein damage before they reach their canonic targets, leading to protein damage response (PDR) in the cells treated. While cells counteract the protein damaging by swiftly damage recognition and clearance, mitochondrial ATP plays a key role in modulating PDR through suppressing protein synthesis mediated by phosphorylation of 4E-BP1 and reducing damaged protein through activating proteasome. We also uncovered a mechanism by which cancer cells, including cancer stem cells (CSCs), evade drug-induced protein damage to acquire MDR, and designed a strategy for overcoming it.

## Results

### The vast majority of anticancer drugs cause protein damage prior to affecting their canonic targets and triggers the protein damage response

We previously showed that 6 of the 11 anticancer drugs tested could induce protein damage (*3*). To provide a more comprehensive understanding of the prevalence of anticancer drugs that can damage proteins and DNA, we first conducted a screen with 101 FDA-approved anticancer drugs in MDA-MB-231 cells for the detection of protein and DNA damage (Supplementary Table 1). The MDA-MB-231 cells were treated with the IC_50_ of each drug for 1 hour and 6 hours, and protein damage was detected on the basis of total protein ubiquitin and DNA damage was detected on the basis of phosphorylation of H_2_AX at Ser139 (*11*), i.e., γH_2_AX (Figure 1A). Our data indicated that at 1 hour post the treatment, 78/101 (77%) and 44/101 (44%) drugs markedly induced protein and/or DNA damage, respectively (Figure 1B, Supplementary Figure 1A, and Supplementary Table 1). We classified the anticancer drugs into 4 groups according to the type of damage that they induced, i.e., drugs that induced both types of damage (BD drugs, 37/101, 37%), damage to proteins only (PD drugs, 41/101, 41%), damage to DNA only (DD drugs, 7/101, 7%), and no damage (ND drugs, 16/101, 16%). We monitored protein aggregation upon 9 anticancer drugs that caused ubiquitination at various levels (Figure 1A), using a molecular rotor dye (PROTEOSTAT) that specifically intercalates into the cross-β spines of the quaternary protein structures found in misfolded and aggregated proteins (*12*). The levels of PROTEOSTAT staining corrected with the levels of ubiquitination very well (Figure 1C), reflecting the correlation of protein structure change with ubiquitination. After 6 hours of treatment, more drugs could induce protein and/or DNA damage (Figure 1B) at higher levels (Supplementary Figure 1B). Notably, at both time points, more anticancer drugs damaged protein than DNA. Similar results were obtained in the A549 lung cancer cell line (Figure 1B, Supplementary Figure 1C, and Supplementary Table 1).

**Fig 1.**
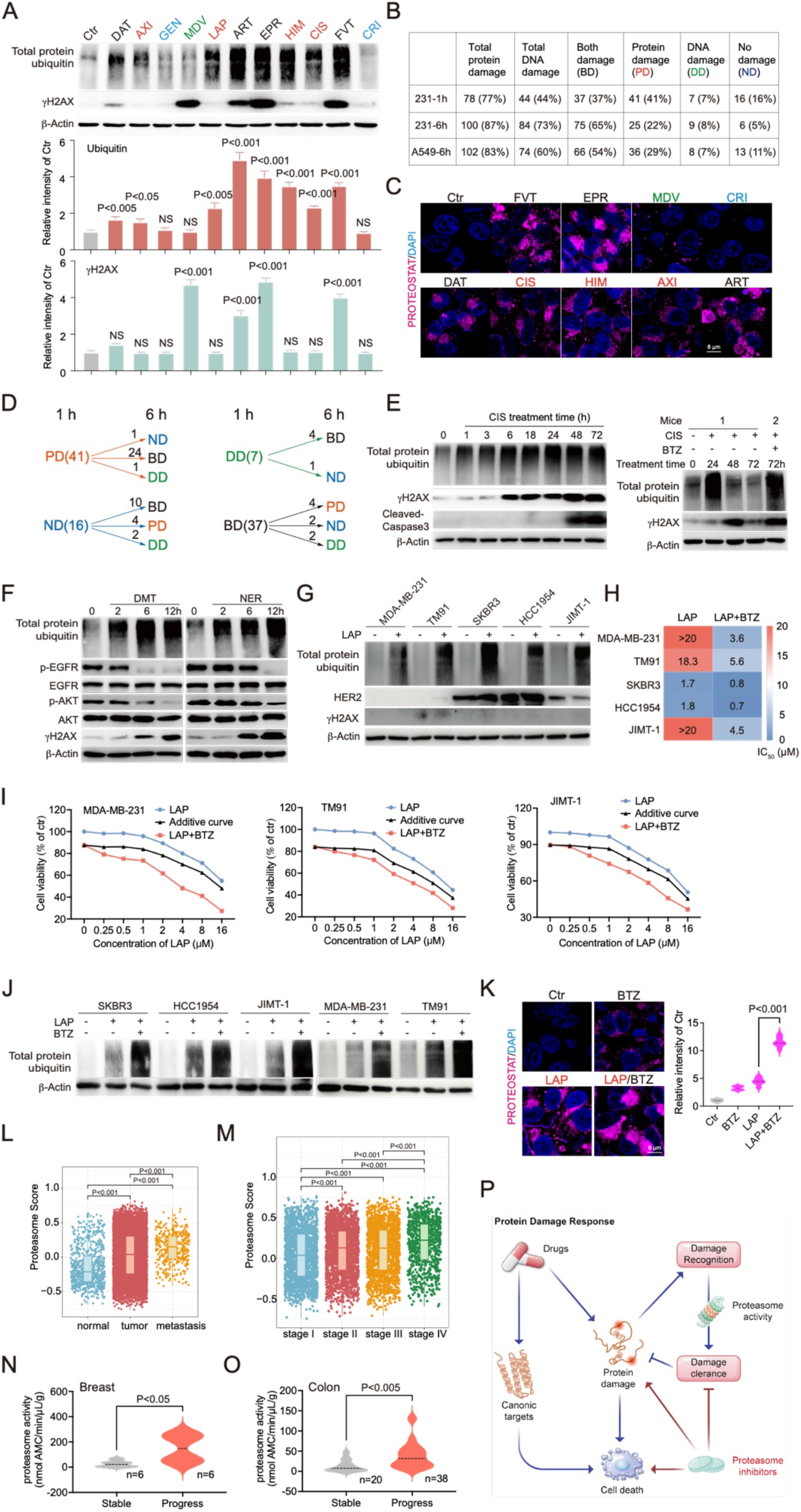
The vast majority of anticancer drugs cause protein damage prior to affect their canonic targets. (A) MDA-MB-231 cells were treated with indicated anticancer drugs for 1 hour, and total protein ubiquitin and γH_2_AX were detected by Western blot. Quantifications of different damages were shown below. (B) Summary of protein and DNA damages induced by anticancer drugs at 1 hour and 6 hours treatment in MDA- MB-231 cells, and 6 hours treatment in A549 cells. Four types of drugs were indicated as BD: both damage, PD: protein damage only, DD: DNA damage only, and ND: no damage drugs. (C) MDA-MB-231 cells were treated with indicated drugs, and protein aggregation were stain by the PROTEOSTAT. (D) Dynamic change of protein damage and DNA damage induced by four types of drugs for 1 hour and 6 hours treatment in MDA-MB-231 cells. (E) MDA-MB-231 cells were treated with *CIS* for indicated time, and total protein ubiquitin, γH_2_AX, and cleaved caspase 3 were detected by Western blot. Nude mice bearing MDA-MB-231 xenograft tumors were treated with vehicle, *CIS* (3 mg/kg) or *BTZ* (1 mg/kg), and total protein ubiquitin and γH_2_AX for tumors were detected at 24 hours, 48 hours, and 72 hours. Tumor tissues were harvested by needle biopsy from mouse 1 at 0, 24 and 48 hours. Both mouse 1 and 2 were sacrificed at 72 hours for obtaining tumors. (F) MDA-MB-231 cells were treated with 2 EGFR inhibitors for indicated time, and total protein ubiquitin, p-EGFR, EGFR, p-AKT, AKT, and γH_2_AX were detected by Western blot. (G) Five breast cancer cell lines were treated with *LAP*, and total protein ubiquitin, HER2, and γH_2_AX were detected by Western blot. (H, I) Five breast cancer cell lines were treated with increasing doses of *LAP* alone or in combination with a single dose of *BTZ*, which achieved 10%-16% killing effect in different cell lines, and cell viability was detected by alamar blue assay. IC50 of *LAP* treatment and *LAP+BTZ* treatment for 5 cell lines were shown (H), and additive curves for 3 *LAP* resistant cell lines were shown (I). Additive = E1+E2-E1*E2, where E1 is inhibition effect from *LAP* and E2 is inhibition effect from *BTZ* treatment. (J) Five breast cancer cell lines were treated with *LAP* alone or in combination with *BTZ* treatment, and total protein ubiquitin were detected by Western blot. (K) MDA-MB- 231 cells were treated with indicated drugs, representative images and quantification of PROTEOSTAT were shown. (L, M) Pan-cancer data from TCGA and TARGET to compare the proteasome activity score (expression of proteasome genes, n=45) in indicated tissue (L) or tumor stages (M). (N, O) The comparision of proteasome activity in resistant patients (PD) and sensitive patients (PR and SD) for the breast cancer patients (n=12) and colon cancer patients (n=58), respectively. The stable and progress disease after chemotherapy were evaluated by the changes of tumor burden according to the RECIST guideline (version 1.1). (P) A summary of protein damage response (PDR). All values are presented as mean value (at least three replications) ± SD, and p value was calculated by comparison with Ctr group or indicated separately.

Next, we studied the dynamics of protein damage and DNA damage induced by 101 drugs that were used to treat MDA-MB-231 cells for both 1 hour and 6 hours (Figure 1D and Supplementary Table 1). We found that 24 out of these 41 drugs moved from the PD group to the BD group, suggesting that although many drugs induced protein damage first, most could damage DNA later. The data also showed that 4 out of 7 drugs moved from the DD group to the BD group; all 16 drugs in the ND group moved to other groups, and a few drugs in the BD group moved to other groups at 6 hours.

Many anticancer drugs have their own well-known primary targets, for example, *cisplatin* (*CIS*) kills cancer cells by damaging DNA through inducing DNA-adducts (*13*). Since our data revealed that protein damage induced by virtually all anticancer drugs occurs quickly upon the treatment, the relationship between the protein damage and the well-known targets of these drugs remains unknow. To investigate this, we first compared dynamic progressions of protein damage and DNA damage induced by *CIS* and the data revealed a time-dependent increase in protein damage starting at 1 hour, and DNA damage appeared several hours later (Figure 1E). To study the prolonged effects of *CIS*, we washed it away after 6 hours of treatment, and we observed increased protein damage at 6 hours, which were cleared away later (Supplementary Figure 2A). While strong DNA damage was observed at 24 hours and lasted for 48 hours, reflecting distinct mechanisms by which cells handle protein damage and DNA damage. In the xenograft tumors, *CIS* induced strong protein damage at 24 hours, and the damaged proteins were gradually cleared away from 24 hours to 72 hours, while DNA damage was observed at 48 hours and repaired at 72 hours (Figure 1E). Meanwhile, proteasome inhibitor (proteasome-I) *bortezomib* (*BTZ*) treatment caused the accumulation of both protein and DNA damage. These results indicate that *CIS* could target the proteins to induce damage prior their canonical target. To further demonstrate this, we tested the *neratinib* (*NER*) and *dacomitinib* (*DMT*), which are selective inhibitors for the epidermal growth factor receptor (EGFR) (*14, 15*). Consistently, our study on two EGFR inhibitors (*NER* and *DMT*) showed that they triggered protein damage at 2 hours, while the inhibition of EGFR and AKT as well as induction of DNA damage became obvious at 6 hours (Figure 1F). Thus, our finding that vast majority of anticancer drugs damage proteins prior reaching their well-known targets may uncover a phenonium, which is rarely recognized, yet occurs commonly upon the treatment of these drugs.

To demonstrate the relationship of anticancer drugs induced protein damage and the drug resistance, we investigated the *Lapatinib (LAP),* a small-molecule tyrosine kinase inhibitor of EGFR and HER2, which has been used for the treatment of HER2-positive breast cancers, however, resistance frequently happens (*16*). When *LAP* was used to treat 2 HER2 negative (MDA-MB-231 and TM91) and 3 HER2 positive (SKBR3, HCC1954 and JIMT-1) breast cancer cell lines (Figure 1G), we found that the HER2 negative cells lines were, in general, more resistant to *LAP* than HER2 positive cells except for JIMT-1 cells, which expresses HER2 at a moderate level and were highly resistant to *LAP* (Figure 1G, H). As *LAP* strongly induces protein damage in these cells (Figure 1G), we tested if blocking the damaged protein clearance by *BTZ* could affect the drug resistance, and the data indicated that the resistance of all 3 cell lines to *LAP* was completely reversed with dramatically increase of protein damage by the *BTZ* treatment (Figure 1H, I, J). While the response of 2 sensitive cell lines to the *BTZ* combination was very miner (Supplementary Figure 2B). Consist with the polyubiquitin level in MDA-MB-231 cells, we tested protein aggregation upon the combination with BTZ, and LAP induced PROTEOSTAT staining was dramatically accumulated by the BTZ treatment (Figure 1K). These data highlight a critical role for protein damage clearance in drug resistance.

To further investigate the relationship between drug resistance and protein damage clearance, we investigated the proteasome activity in the pan-cancer dataset encompassing over 10,000 samples spanning 40 cancer types, which were aggregated from The Cancer Genome Atlas (TCGA) and Therapeutically Applicable Research to Generate Effective Treatments (TARGET) databases. The data indicated that the proteasome activity was higher in the tumor and further increased in the more malignant metastasis (Figure 1L), implying that the proteasome activity is associated with the tumor malignancy. Indeed, with the develop of tumor stage, the proteasome activity gradually increased in the pan-solid tumor (Figure 1M). We then followed 12 breast patients who received various drug treatment for their responses to the drug treatment and proteasome activity (Supplementary Figure 2C). Our data indicated significant higher proteasome activity in patients who exhibited disease progression (PD, n=6) than patients who displayed partial response (PR, n=5) or stable disease (SD, n=1) (Figure 1N). Our data on 58 colon cancer patients, who received various drug treatment regimen, also detected higher proteasome activity in 38 patients who developed PD that 20 patients with PR or SD (Figure 1O and Supplementary Figure 2D). Consistently, in the TCGA and GES databases, two cohorts of colon cancer patients, which treated with Capecitabine and Bevacizumab, respectively, the resistant patients (non-response) have significant higher proteasome activity score compared to the sensitive patients (response), and two other types of cancer were also identified to have higher proteasome activity score in the resistant patients (Supplementary Figure 2E).

Altogether, these results suggest a model through which majority of anticancer drugs damage proteins prior to reaching their canonic targets, triggering protein damage response (PDR). The PDR contains two major steps, i.e. the damage recognition, which is mainly mediated by ubiquitin system and the damage clearance, which is mainly mediated by proteasome system as the treatment of proteasome inhibitors could result in the accumulation of protein damage, leading to the accelerated cell death (Figure 1P). Conversely, cells with higher levels of proteasome activity intrinsically or gradually forced to increase proteasome activity should survive better to the cytotoxicity of anticancer drugs, which might be a reason why the drug resistant breast and colon cancers have higher levels of proteasome activity than the drug responders (Figure 1N,O).

### Anticancer drugs damage neosynthesized proteins and impair their functions

Next, we attempted to understand the mechanism underlying protein damage induced by anticancer drugs, we used various inhibitors with anticancer drug treatment. While cycloheximide (CHX), which blocks protein synthesis on the ribosome (*17*), completely blocked total protein ubiquitin induced by the three anticancer drugs tested (*genistein*, *GEN*; *artemether*, *ART*; and *lapatinib*, *LAP*), whereas no obvious effects were observed upon treatment with all other factors tested, including salubrinal (Salu), which inhibits endoplasmic reticulum (ER) stress; N-acetyl-l-cysteine (NAC) and L-ascorbic acid (VC), which block ROS accumulation; and thymidine (dThd), which blocks the cell cycle progression (*18*) (Figure 2A). These data suggest that protein synthesis is the main target of anticancer drug-induced protein damage, and the data also revealed that the anticancer drugs had distinct effects, causing protein damage and DNA damage. For example, CHX blocked protein damage and induced DNA damage, while dThd induced DNA damage but did not induce protein damage (Figure 2A).

**Fig 2.**
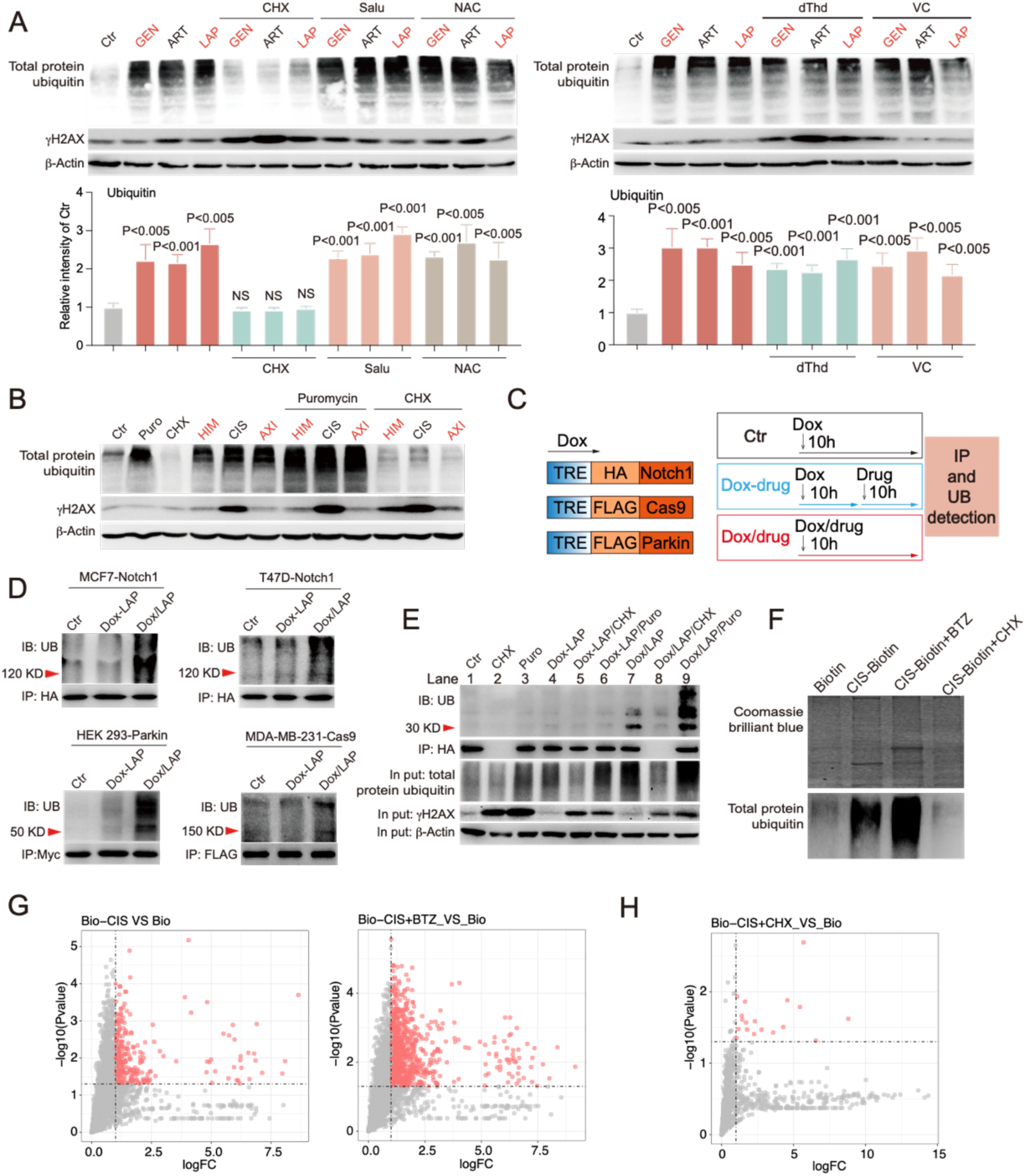
Anticancer drugs damage neosynthesized proteins and impair their functions. (A) MDA-MB-231 cells were treated with indicated anticancer drugs or combined with various inhibitors, respectively. Total protein ubiquitin and γH_2_AX were detected by Western blot and quantification for ubiquitin was shown below. (B) MDA-MB-231 cells were treated with indicated anticancer drugs or combined with CHX or puromycin, and total protein ubiquitin and γH_2_AX were detected by Western blot. (C) Treatment model for comparation the damages between matured protein and neosynthesized proteins. (D) MCF7, T47D, HEK293, and MDA-MB-231 cells transfected with doxycycline inducible Notch1, Parkin, or Cas9 respectively. Cells were treated with *LAP* as indicated strategy, and Notch1, Parkin, or Cas9 were pulled down for detection of ubiquitination. (E) MDA-MB-231-GFP cells were treated with *LAP* or combined with protein synthesis inhibitors, and GFP were pulled down for detection of ubiquitination, and total protein ubiquitin and γH_2_AX was detected by Western blot. (F-H) MDA-MB-231 cells were treated with indicated drugs, and biotin was pulled down, and interacting proteins were stained with Coomassie brilliant blue and protein ubiquitin were detected by Western blot (F). *CIS*-biotin and biotin interacting proteins were analyzed by mass spectrum and enhanced binding proteins were highlight by log_2_ FC>1, P value<0.05 (G, H). All values are presented as mean value (at least three replications) ± SD, and p value was calculated by comparison with Ctr group or indicated separately.

To further understand the role of protein synthesis in drug-induced protein damage, we compared the effects of CHX, which completely blocks protein synthesis, and puromycin, a ribosome inhibitor that interferes with protein synthesis via its incorporation into the C-terminus of nascent polypeptides, which leads to premature termination of peptide synthesis (*19*). Strikingly unlike CHX treatment, which completely blocked protein damage induced by all drugs, puromycin treatment failed to block protein damage and instead it significantly enhanced protein damage induced by the drugs (Figure 2B). Because puromycin treatment generated more prematurely terminated nascent peptides, this may account for the significant increase in ubiquitination. Altogether, our data suggest that newly synthesized proteins and nascent peptides might be the main targets of anticancer drug-induced damage.

To further investigate this hypothesis, we applied the doxycycline-inducible Tet-on system to investigate protein damage during and after the synthesis of several specifically designed proteins using the protocol shown in Figure 2C. In the first group (Ctr), the expression of the tested genes was induced by doxycycline treatment for 10 hours, but no drugs were used. In the second group (dox-drug), doxycycline was added to induce protein synthesis for 10 hours, after which the doxycycline was washed away to stop protein synthesis prior to *LAP* treatment for another 10 hours. In the third group (dox/drug), doxycycline and the *LAP* were added together to expose newly synthesized proteins to *LAP* treatment. The test proteins were then pulled down by immunoprecipitation, and their ubiquitination was detected (Figure 2C). For all three proteins, HA-Notch1, Myc-Parkin and FLAG-Cas9, tested in 4 cell lines, MCF7, T47D, HEK293, and MDA-MB-231 cells, the results consistently demonstrated that protein damage occurred in the 3^rd^ group (Figure 2D), i.e., the anticancer drugs preferably damaged newly synthesized proteins but had very little effect on the same proteins that were already synthesized. To further demonstrate this, we interfered with target protein synthesis by CHX or puromycin (Figure 2E). The *LAP* treatment-induced ubiquitination of newly synthesized GFP (lane 7) was completely blocked by the inhibition of GFP synthesis (CHX, lane 8) and further increased by the premature termination of GFP synthesis (Puromycin, lane 9). Comparison of Dox-*LAP*/Puro (lane 6) and Dox/*LAP*/Puro (lane 9) treatment clearly demonstrated that *LAP*/Puro treatment caused a strong increase in total protein damage, while the damage to GFP occurred only in synthesizing proteins (lane 9) and not in already synthesized or matured GFP proteins (lane 6). *LAP* is a small-molecule tyrosine kinase inhibitor of EGFR and HER2, the damage of Notch1, Parkin, Cas9, and GFP indicate that anticancer drugs could damage the proteins nonspecifically which mainly comes from the newly synthesized proteins and nascent peptides.

We suspected that these drugs might be able to bind proteins during or shortly after their synthesis for causing the damaging. To investigate this, we covalently linked *CIS* to biotin to generate the *CIS*-biotin and used it to treat MDA-MB-231 cells for 3 hours for generate protein damaging as revealed by their ubiquitination, and then, *CIS*-biotin was pulled down by the streptavidin immobilized on agarose and visualized by Coomassie brilliant blue staining (Figure 2F). The results indicated that *CIS*-biotin pulled down more damaged proteins compared to the biotin alone, and the damaged proteins were further increased when combined with *BTZ* treatment and were decreased by CHX treatment as reflected by intensity of protein ubiquitination (Figure 2F). Mass spectrometry analysis identified 287 proteins (log_2_ FC>1, P value<0.05) pulled down by the *CIS*-biotin after extracting proteins bound by biotin alone, and the number increased to 865 (log_2_ FC>1, P value<0.05) when the damaged protein clearance was blocked by the *BTZ* treatment whereas the number of proteins reduced to 18 (log_2_ FC>1, P value<0.05) upon CHX treatment (Figure 2G, H). Because the large quantity of proteins is involved with a short time window, we believed that the drug might bind to proteins nonspecifically. Thus, these results, in principle, indicate that anticancer drugs might bind and damage proteins nonspecifically before reaching their canonical targets.

Next, we asked whether the protein damage induced by anticancer drugs could affect protein function. Since ubiquitination levels of the newly synthesized proteins were higher, we tested the function of these proteins by investigating GFP fluorescence intensity and luciferase activity. MDA-MB-231-GFP cells were treated with 6 drugs during GFP synthesis (dox/drug) or after GFP synthesis (dox-drug). GFP fluorescence was dramatically decreased in the dox/drug group compared to the dox-drug group (Supplementary Figure 3A, B). There was no significant change in the GFP level in these groups (Supplementary Figure 3C), suggesting that the decline in GFP fluorescence was caused by protein damage and not by changes in protein abundance. Similar damage to synthesizing luciferase was induced by 5 structurally unrelated drugs (Supplementary Figure 3D and E). Altogether, our data demonstrate that anticancer drugs interact nonspecifically with newly synthesized proteins and impair their function rather than affecting their synthesis.

### Protein damage occurs at low levels in quiescent cells but high levels in proliferating cells

Quiescence, dormancy and slow cycling are key features of many drug-resistant models, including CSCs, cells under hypoxia, and persistent cancer cells, and the survival of these slow-growing cells under chemotherapy, radiotherapy, and immune checkpoint blockade therapy leads to drug resistance (*20–22*). However, the underlying mechanism is unknown. Our analysis thus far identified neosynthesized proteins as the main targets for damage and that cell proliferation places a high demand on protein synthesis (*23*). To investigate this mechanism, we first compared protein damage in quiescent cells generated by 3 days of serum starvation and actively proliferating cells grown in regular culture containing 10% serum. We found that the quiescent cells almost completely evaded drug-induced protein damage, reproducing the effect of CHX treatment, while the proliferating cells accumulated extensive damaged proteins (Figure 3A). We noticed that the quiescent cells also evaded drug-induced DNA damage. To study drug-induced cell death, we monitored the dynamics of caspase 3 activation using a fluorescence resonance energy transfer (FRET)[based caspase 3 (C3) biosensor system (*24*) (Supplementary Figure 4A). Indeed, cell death induced by the anticancer drugs was dramatically decreased in the quiescent cells compared to the proliferating cells (Figure 3B and C), which is consistent with the levels of protein damage induced by the drugs (Figure 3A), indicating that the evasion of protein damage is a key mechanism leading to MDR.

**Fig 3.**
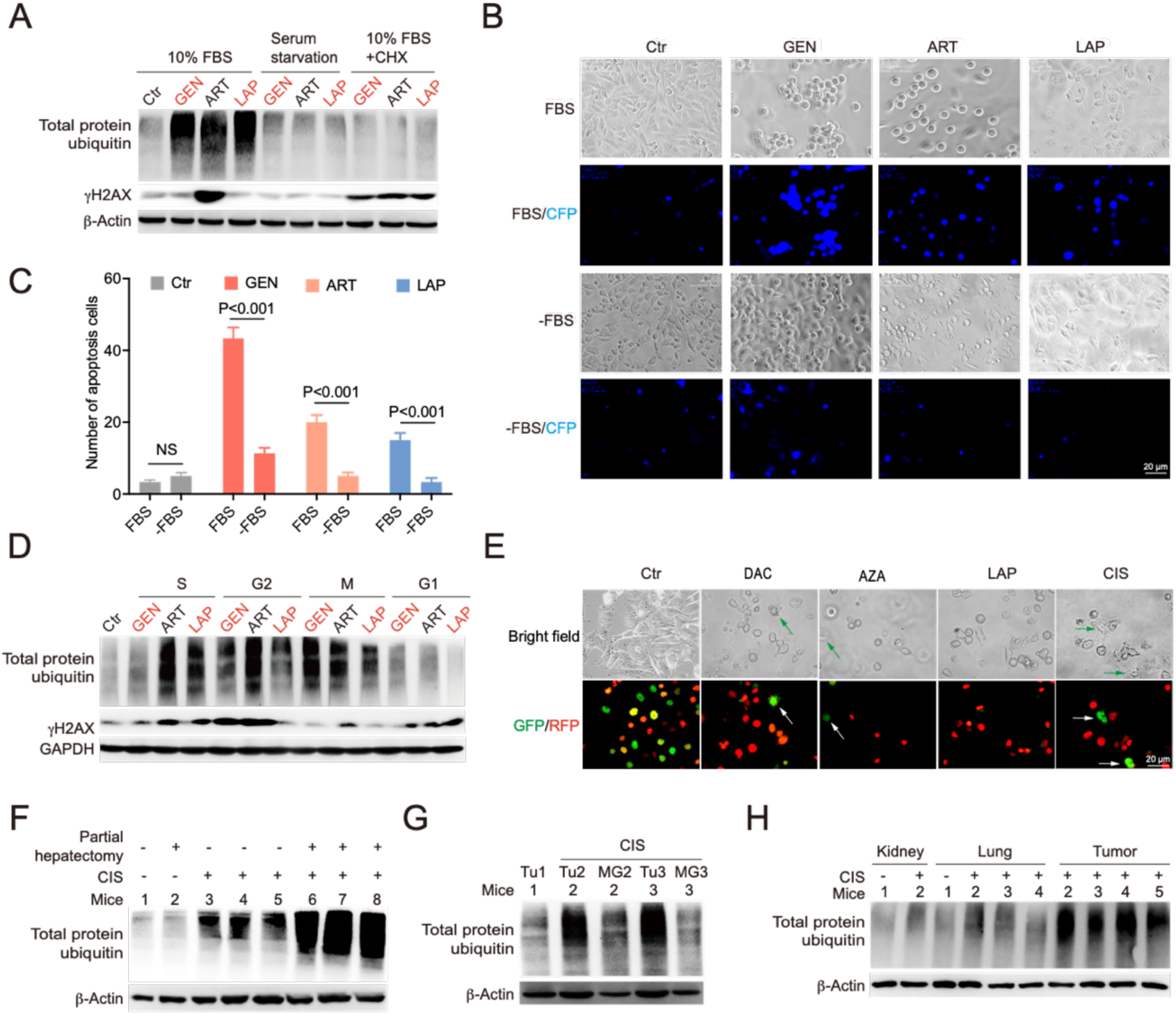
Protein damage occurs at low levels in quiescent cells but high levels in proliferating cells. (A) MCF7 cells under 10% FBS culture or serum starvation for 3 days were treated with indicated anticancer drugs and total protein ubiquitin and γH_2_AX were detected by Western blot. (B, C) MDA-MB-231-C3 cells were under 10% FBS culture or serum starvation for 3 days and then treated with indicated drugs, and caspase 3 activation was monitored by CFP fluorescence (B) and quantified (C). (D) MDA-MB-231 cells were synchronized by double thymidine treatment, and cells at S phase, G2 phase, M phase, or G1/G0 phase were treated with indicated drugs. Protein ubiquitin and γH_2_AX were detected by Western blot. (E) Illustration of changes of MDA-MB- 231-CG cells, which are labeled with mKO2-Cdt1 at S/G2/M phases (green) and mAG-Geminin at G1/G0 phases (red), upon the treatment of anticancer drugs as indicated for 7 days. (F) Eight 5-month-old mice with or without partial hepatectomy were treated with *CIS* (6 mg/kg) for 24 hours, and total protein ubiquitin was detected by Western blot. (G, H) BALB/c mice carrying 4T1 allograft tumors were treated with *CIS* (6 mg/kg) for 24 hours, and total protein ubiquitin for tumors, adjacent normal tissue, and several organs were detected by Western blot. All values are presented as mean value (at least three replications) ± SD, and p value was calculated by comparison with Ctr group or indicated separately.

To further understand the relationship between protein damage and cell proliferation, we released cells into S, G2, M, and G1 phases at various time points after synchronization of the cells by double thymidine block (Supplementary Figure 4B) and measured their protein damage levels. Although some variations depending on cell cycle phase existed, the cells at proliferative phases (S, G2, and M) accumulated more protein damage than cells in G1 phase (Figure 3D). To further demonstrate this, we used MDA-MB-231-CG cells that were labelled with mKO2-Cdt1 (G1/G0 phase, red) and mAG-Geminin (S/G2/M phases, green) to visualize the cell cycle (Supplementary Figure 3C) (*25*). The data indicate that a higher percentage of S/G2/M cells were associated with enhanced protein damage induced by anticancer drugs (Supplementary Figure 4C and D). The proliferative (green) cells were virtually eradicated, whereas nonproliferating (red) cells survived the treatment of 5 structurally unrelated drugs, respectively (Figure 3E). In summary, our results indicate that drug-induced protein damage mainly occurs during the proliferative phases (S/G2/M) and that cells at non-proliferative phases (G0/G1) accumulate much less protein damage.

To validate our discovery in vivo, we first compared protein damage under *CIS* treatment in fast-proliferating liver (induced by partial hepatectomy, Supplementary Figure 4E) and intact adult mouse liver (*26*), and indeed, protein damage in the fast-proliferating liver tissue was more pronounced than that in the normal mouse liver tissue (Figure 3F). It is well known that tumors undergo faster proliferation than adjacent normal tissues (*27*). Thus, we treated BALB/c mice bearing 4T1 tumors with *CIS*, and the level of protein damage in the tumor cells was much stronger than that in the adjacent mammary tissues (Figure 3G). Cells from the kidney, lung, gut, heart, and spleen were also analyzed, and similarly, *CIS* caused much less protein damage in these normal tissues than in the tumors (Figure 3H, Supplementary Figure 4F). Altogether, our data indicate that protein damage occurs at low levels in quiescent cells but high levels in proliferating cells.

### ATP positively regulates protein synthesis and dictates sensitivity to anticancer drugs

Next, we investigated the mechanism underlying differential protein damage in quiescent cells and proliferating cells. The reduction in damaged protein accumulation in quiescent cells can be due to decreased damaged protein loading, enhanced damaged protein clearance by the proteasome, or a combination of both. To distinguish between these scenarios, we first examined protein synthesis, which contributes to damaged protein loading (Figure 2D). Quiescent cells and cells at different phases of the cell cycle were treated with puromycin, which is incorporated into nascent chains during protein synthesis and serves as a marker for protein synthesis. The results indicated that quiescent cells and cells at G1 phase had significantly lower levels of protein synthesis than proliferating cells (Figure 4A). Protein synthesis is strictly regulated by metabolism (*28, 29*) and consumes the largest proportion of ATP. Thus, we tested whether ATP could affect protein synthesis. We first checked whether ATP could enter the cells. ATP was incubated with parental MDA-MB-231 cells and MDA-MB-231-R3 (231-R3) cells in a multidrug-resistant state that we previously generated (*3*). Our data showed a time- and dose-dependent increase in the ATP level inside the cells (Figure 4B). Associated with this was an increase in protein synthesis in both the MDA-MB-231 cells and 231-R3 cells (Figure 4C). We showed earlier that quiescent cells could evade anticancer drug-induced cell apoptosis (Figure 3B). We suspected that this might be due to their low levels of ATP. To demonstrate this, we treated quiescent cells (under serum starvation culture conditions) with ATP and found that ATP treatment indeed dramatically increased the protein synthesis (Figure 4D). Associated with this increase in protein synthesis was the enhanced protein damage (Figure 4E) and cell apoptosis (Figure 4F) under *CIS* treatment in quiescent cells. In summary, these results indicate that ATP enhances protein synthesis, which contributes to the damaged protein loading and increased the cytotoxicity of anticancer drugs.

**Fig 4.**
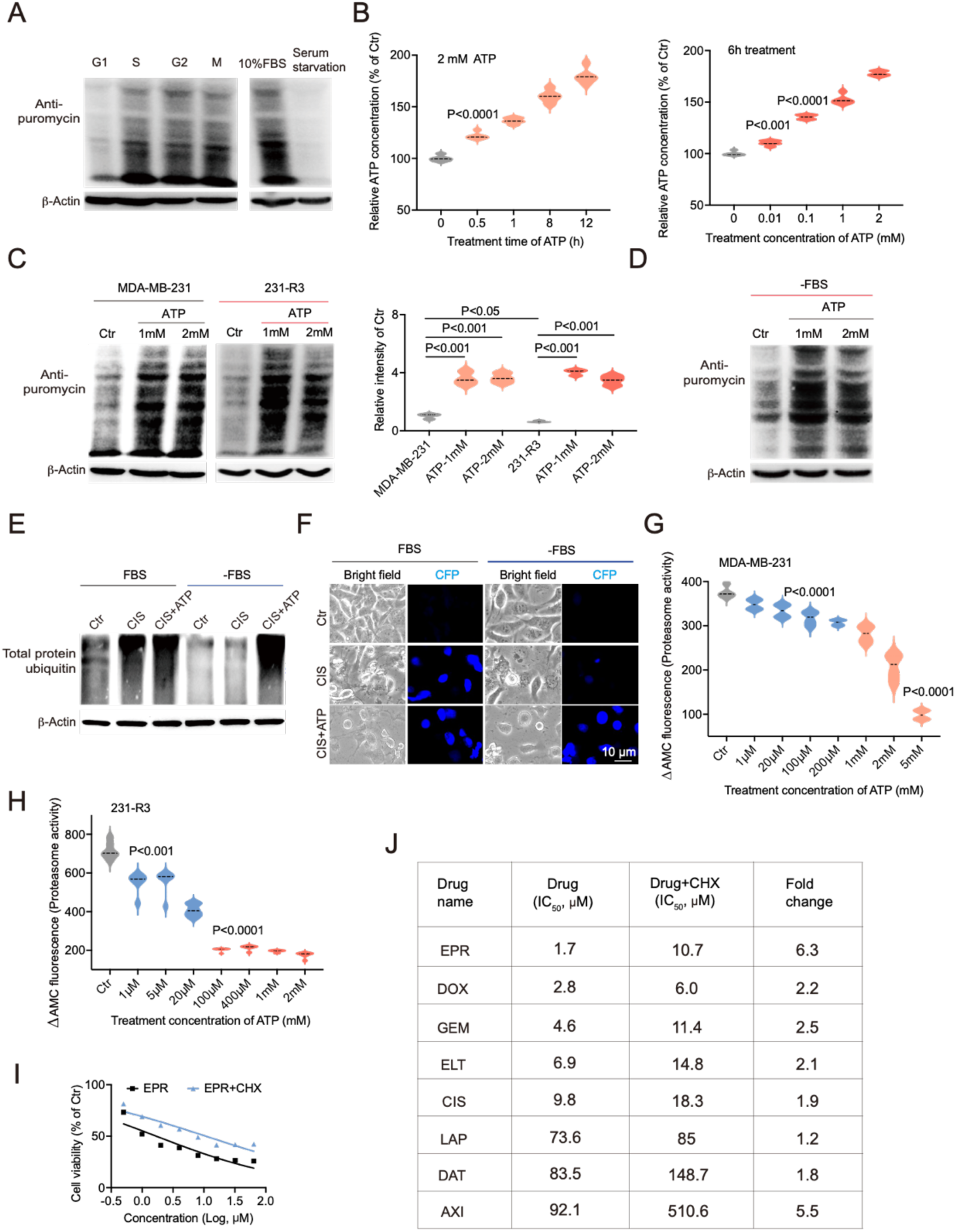
ATP positively regulates protein synthesis and dictates sensitivity to anticancer drugs. (A) MDA-MB-231 cells at various conditions were treated with 10mg/mL puromycin for 15 min, and protein synthesis was measured by Western blot using an antibody to puromycin. (B) Relative ATP quantity inside cells over time or upon varying doses of ATP treatment were detected by an ATP detection kit. (C) 231-R3 and MDA-MB-231 cells were treated with 1- or 2-mM ATP, and protein synthesis were measured by Western blot and quantified. (D) MDA-MB-231 cells under quiescent state (FBS starvation) were treated with ATP, and protein synthesis were measured by Western blot. (E) MDA-MB-231-C3 cells under 10% FBS culture or quiescent state (FBS starvation) were treated with *CIS* or combined with 2 mM ATP, and protein damage was detected by Western blot. (F) MDA-MB-231-C3 cells under 10% FBS culture or quiescent state were treated with *CIS* or combined with 2 mM ATP, and caspase 3 activation was monitored by the CFP fluorescence. (G, H) MDA-MB-231 and 231-R3 cells were treated with increasing dose of ATP, and the proteasome activity were measured by fluorescent AMC tagged peptide substrate. Please note that 231-R3 cells have higher levels of proteasome activity and are more responsive to ATP. (I, J) MDA-MB-231 cells were treated with indicated anticancer drugs or combined with CHX treatment, and cell viability was detected by Alamar Blue assay (I) and IC_50_ was fitted by Prism-GraphPad-9.0.0 (J). All values are presented as mean value (at least three replications) ± SD, and p value was calculated by comparison with Ctr group or indicated separately.

ATP production at a physiological level is a negative regulator of proteasome activity (*30, 31*). We have previously shown that the 231-R3 cell line produces less ATP and has higher proteasome activity than the parental MD-MBA-231 cells (*3*). To investigate the potential impact of ATP in these cells, we treated them with ATP and found that it could increase protein synthesis in both cell lines (Figure 4C). Meanwhile, a dose-dependent decrease in the proteasome activity was found in both cell lines, with a sharper reduction in proteasome activity in the 231-R3 cells (Figure 4G, H), which might be due to the initial lower ATP level and higher proteasome activity in the 231-R3 cells. Collectively, our data demonstrate that ATP plays a positive role in protein synthesis and a negative role in proteasome activity, with both roles affecting the levels of protein damage and responses to anticancer drugs.

We have shown earlier that inhibition of the protein synthesis could block the protein damage, yet the relationship between protein synthesis and MDR had not been tested. To study this, we treated fast-proliferating MDA-MB-231 cells with increasing concentrations of drugs in the absence or presence of CHX, which blocks protein synthesis. The data revealed that all 8 drugs tested exhibited increased IC_50_ values to varying degrees, ranging from 1.2-fold to 6.3-fold higher in the presence of CHX (Figure 4I, J, and Supplementary Figure 5), indicating that the reduction in protein synthesis resulted in increased MDR.

### Maintenance of a low-ATP status is a common strategy by which CSCs and drug-resistant acquired cells (DRACs) acquire MDR

ATP production by mitochondrial oxidative phosphorylation (OXPHOS) exceeds glycolytic ATP generation by at least 14.5-fold (*32*). Since ATP production dictates the ability of anticancer drugs to damage proteins, mitochondrial OXPHOS should be the key factor that regulates drug-induced protein damage in quiescent cells and proliferative cells. Indeed, we found that the quiescent cells had a much lower mitochondrial oxygen consumption rate (OCR), mitochondrial ATP production, and total ATP level than the cells under regular growth conditions (Figure 5A and Supplementary Figure 6A). On the other hand, we detected the mitochondrial OCR and ATP production in the proliferative cells and G1 cells, and the results indicated that the mitochondrial OCR was significantly increased in the proliferative cells (S/G2/M cells) compared to the G1 cells, and ATP production also showed a similar trend (Figure 5B). Thus, the mitochondrial OCR is tightly associated with drug-induced protein damage in quiescent cells and proliferative cells revealed earlier (Figure 3A and D). To further validate that the mitochondrial OCR regulates protein damage, *LAP* was combined with antimycin A and oligomycin treatment and the data indicated that the treatment suppressed the mitochondrial OCR and ATP production (Supplementary Figure 6B), and markedly decreased protein damage in all phases of the cell cycle (Figure 5C). Altogether, these findings demonstrate that the mitochondrial OCR, which linked to ATP, played a critical role in the drug induced protein damage.

**Fig 5.**
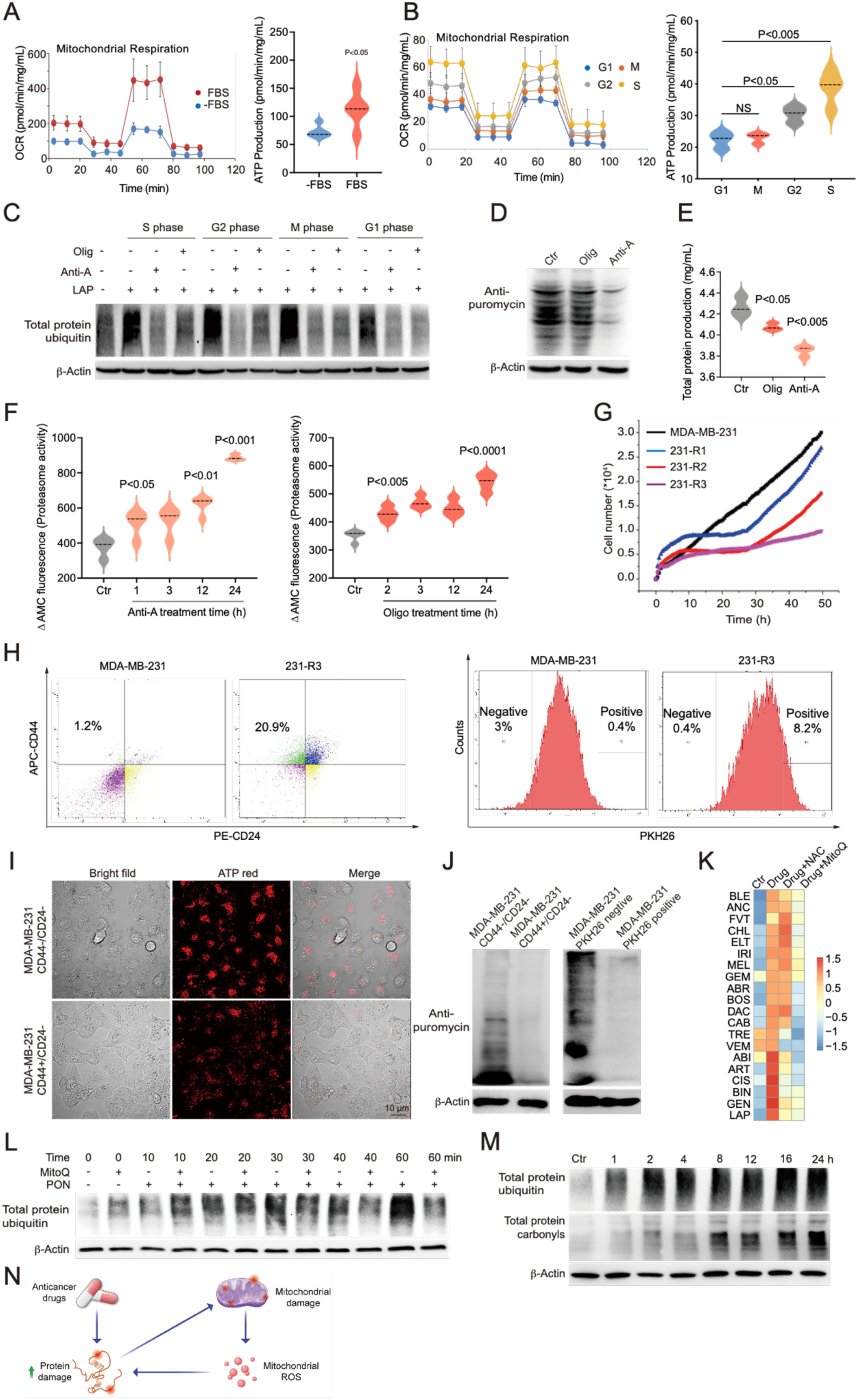
Maintenance of a low-ATP status is a common strategy by which CSCs and drug-resistant acquired cells (DRACs) acquire MDR. (A and B) Mitochondrial OXPHOS and ATP production were detected by Seahorse XFp Cell Mito Stress Test assays for the blow cells: MDA-MB-231 cells at S phase, G2 phase, M phase, G1 phase, MDA-MB-231 cells under 10% FBS culture and quiescent state (FBS starvation). (C) MDA-MB-231 cells at different cell cycle were treated with *LAP* or combined with antimycin A or oligomycin treatment, and total protein ubiquitin was detected by Western blot. (D, E) MDA-MB-231 cells were treated with oligomycin or antimycin A, and protein synthesis were measured by Western blot (D), and total protein production was detected by BCA protein assay kit (E). (F) MDA-MB-231 cells were treated with oligomycin or antimycin A for indicated time, and proteasome activity were measured by fluorescent AMC tagged peptide substrate. (G) Cell proliferation rate of MDA-MB-231, 231-R1, 231-R2, and 231-R3 cells. (H) MDA-MB-231 and 231-R3 cells were labeled with APC-CD44 and PE-CD24, and CD44+/CD24- and CD44-/CD24- cells were sorted out. MDA-MB-231 and 231-R3 cells were labeled with PKH26 and cultured for 2 weeks, and PKH26 high and PKH26 low cells were sorted out. (I) CD44+/CD24- and CD44-/CD24- MDA-MB-231 cells were stained with a cell-permeable red fluorescent probe for the ATP. (J) CD44+/CD24-, CD44-/CD24-, PKH26 high, and PKH26 low MDA-MB-231 cells were labeled with puromycin, and protein synthesis was detected by Western blot. (K) heatmap for the intensity of total protein ubiquitin by single drug treatment or combined with NAC, MitoQ treatment (n=20). (L) MDA-MB-231 cells were treated with ponatinib alone or combined with MitoQ for indicated time, and protein total ubiquitination was detected by western blot. (M) MDA-MB-231 cells were treated with ponatinib for indicated time, and protein total ubiquitination and protein total carbonylation were detected by western blot. (N) A summary of interplay between anticancer drugs and mitochondrial ROS to protein damage. All values are presented as mean value (at least three replications) ± SD, and p value was calculated by comparison with Ctr group or indicated separately.

Because we showed earlier that ATP could affect the protein synthesis and proteasome activity, we hypothesized that the mitochondrial OCR might regulate both. To investigate this possibility, MDA-MB-231 cells were treated with oligomycin or antimycin A, both of which significantly reduced the protein synthesis (Figure 5D), which was accompanied by a decrease in the total protein content (Figure 5E). At the same time, we measured the proteasome activity and found that oligomycin or antimycin A treatment led to a time-dependent increase in the proteasome activity (Figure 5F). As expected, proliferative cells at all phases had decreased proteasome activity compared to that of G1 phase cells and Ctr cells, which were a mixture of cells at all phases of the cell cycle (Supplementary Figure 6C). Collectively, these results indicate that quiescent cells have a minimal mitochondrial OCR and ATP production, which is critical to decrease damaged protein loading and enhances damaged protein clearance, both of which resulted in the evasion of drug induced protein damage.

We previously generated the 231-R1, 231-R2, and 231-R3 drug-resistant acquired cells (DRACs), which acquired increasing drug resistance during long-term exposure to gradually increasing doses of *CIS* (i.e., 1 μM, 3.5 μM, and 10 μM) (*3*). We found that the resistance of DRACs was negatively correlated with their growth rate, with the most drug resistant 231-R3 cells exhibiting the lowest growth rate (Figure 5G), indicating that the DRACs might also have decreased metabolic flux. Indeed, our previous data demonstrated that ATP production gradually declined in 231-R1, 231-R2, and 231-R3 cells (*3*), indicating that the maintenance of low-ATP level might also be the strategy for DRACs to acquire MDR. Notably, we knocked down the key genes in mitochondrial OXPHOS (NDUFS1 and NDUFA2) in the parental MDA-MB-231 cells, which markedly decreased the cell proliferation rate (Supplementary Figure 6D), accompanied by the resistance to *CIS* (*3*). Collectively, these studies show that the reduction in mitochondrial OXPHOS to reduce ATP production is a strategy by which DRACs to develop the MDR.

CSCs are defined by their fundamental quiescent nature, which is believed to be the primary cause of drug resistance in multiple types of cancers (*33*). To investigate if the mechanism of drug resistance identified here is applicable to CSCs, we sorted MDA-MB-231 and 231-R3 cells using CD44+/CD24- expression, which is commonly used to identify breast CSCs (*34*). The data detected 20.9% and 1.2% CD44+/CD24- cells in the 231-R3 cell line and MDA-MB-231 cell line, respectively (Figure 5H). Similarly, using the dye PKH26, which tracks stem cells as its accumulation is higher in cells undergoing slower division (*35*), we detected 20-fold higher levels of PKH26+ cells in 231-R3 cells than in MDA-MB-231 cells (Figure 5H). These findings are consistent with the slower proliferation rate and higher levels of drug resistance of the 231-R3 cells compared to MDA-MB-231 cells and suggesting that CSCs might also adopt the same strategy to develop the MDR. Thus, we compared the levels of ATP and protein synthesis in the CD44+/CD24- cells and CD44-/CD24- cells isolated from MDA-MB-231 cells, and the data revealed significantly lower levels of ATP and protein synthesis in the CD44+/CD24- cells than in the CD44-/CD24- cells (Figure 5I, J). We also observed a significant decrease in protein synthesis in the PKH26-positive MDA-MB-231 cells (Figure 5J). Collectively, our data indicate that the maintenance of low ATP levels is a common strategy by which CSCs and DRACs to acquire MDR.

Next, we investigated the mechanism underlining the decreased mitochondrial OXPHOS in drug resistance cells. In our earlier data, we demonstrated the cytosol ROS is not involved in the protein damage of 3 tested drugs (Figure 2A), while we tested the protein carbonylation, which is the marker of protein oxidative damage (*36*), 9 out of 11 drugs induced significant protein oxidation (Supplementary Figure 6E). The cytosol ROS and mitochondrial ROS have distinct physiology (*37, 38*), so we compared the effect of inhibitors for cytosol ROS and mitochondrial ROS. Notably, we found MitoQ, which is a mitochondrial ROS inhibitor, was the most potent in blocking drug induced protein damage compared with other 5 cytosol ROS inhibitors (Supplementary Figure 6F) and could attenuate the protein damage for all 20 drugs tested while NAC could only mildly attenuated protein damage for 8 out of the 20 drugs (Figure 5K, Supplementary Figure 6G and Supplementary Figure 7C). These results indicate that mitochondrial ROS might be commonly induced by anticancer drugs. Indeed, we detected 3 out of 11 drugs could generate the cytosol ROS (Supplementary Figure 7A), and all of the 11 drugs could induce the mitochondrial ROS accumulation (Supplementary Figure 7B). These data suggested that mitochondrial ROS is commonly generated and contributes to drug induced protein damage. Since our earlier finding indicates that ribosome mediated protein neo-synthesis is the major source of drug induced protein damage, we compared MitoQ and CHX treatment, and the data indicated that MitoQ only partially blocked the protein ubiquitination whereas CHX treatment completely blocked the damage (Supplementary Figure 7C), indicating that damage of the neosynthesized protein might occur first. To validate this, we tested several drugs, such as ponatinib (PON) and found that PON induced protein damage starting at 10 min and reached a pick at 60 min and the effect of MitoQ on protein damage was not observed until 30 min later and became strong at 60 min (Figure 5L). Our data also showed that PON treatment induced mitochondrial ROS was not observed at 1 hour and became obvious starting from 8 hour (Supplementary Figure 7D). Thus, the drug treatment might damage the neosynthesized protein directly prior to mitochondrial ROS. To demonstrate this, we used the puromycin treatment to generate the noepeptide (*19*) followed by incubation with or without PON treatment, and the data showed that the ubiquitination of nascent peptides was significantly increased by PON treatment (Supplementary Figure 7E). The damaged neopeptide should contain mitochondrial proteins, since 99% of them is synthesized in the cytosol ribosomes (*39*), and the damaged mitochondrial proteins might cause the generation of mitochondrial ROS. Indeed, increase of protein damage through BTZ treatment significantly enhanced PON induced mitochondrial ROS, while the blockage of neoprotein synthesis by CHX treatment eliminate the PON induced mitochondrial ROS (Supplementary Figure 7D). These results indicate that the anticancer drugs induce neopeptide damage first, which contribute to the mitochondrial ROS accumulation, leading to more extensive oxidative protein damage. Indeed, we observed the PON treatment induced protein ubiquitination at 1 h, and the protein carbonylation were detected at 8 hours (Figure 5M). Collectively, these data suggest a positive feedback loop between anticancer drugs and mitochondrial ROS in such a way that the anticancer drugs damage the neo peptide at early phase of drug treatment, which results in the mitochondrial damage, and increased mitochondrial ROS generation, leading to the enhanced protein damaging (Figure 5N). Thus, maintenance of a low mitochondrial activity gives CSCs and DRACs the advantage to escape the early phase of protein damage by decreasing ATP production, which reduce the neo peptide synthesize, and later phase of protein damage by decreasing the mitochondrial ROS generation, which might damage the neo peptide directly.

### PDK1 inhibition forces ATP production that upregulates protein synthesis by enhancing p4E-BP1

Our data thus far demonstrated that decreased ATP production due to reduced mitochondrial OXPHOS plays a central role in drug-induced protein damage and MDR. To reverse the low-ATP strategy observed in the cells with intrinsic and acquired resistance, we analyzed the expression of mitochondrial-related genes and found that some genes were upregulated in the 231-R3 cells (Figure 6A). Among these genes, pyruvate dehydrogenase kinase 1 (PDK1) was the most significantly activated, together with PDHX and LDHAL6A, which encode downstream enzymes. PDK1 regulates the metabolic shift from OXPHOS to glycolysis, known as the Warburg effect (*40*). It was recently shown that although cancer cells use more ATP generated by glycolysis than normal cells, over 80% of the ATP that they use is still derived from mitochondrial OXPHOS (*32*). Thus, PDK1 overexpression, which inhibits mitochondrial ATP production, might elicit a profound effect on cancer cells. We found that PDK1 expression was higher in the 231-R3 cells, and it was further enhanced after treatment with *CIS* (Figure 6B), which is consistent with the decreased mitochondrial ATP production in 231-R3 cells (*3*). To reverse the low-ATP strategy used by resistant cells, we decided to target PDK1. We first tested 2 PDKI-Is, dichloroacetophenone (DAP) and P64, which is a DAP derivative with improved selectivity and antiproliferative activity (*41*). Treatment with either drug significantly enhanced the mitochondrial OCR, mitochondrial ATP level, and total ATP level (Figure 6C and Supplementary Figure 8A, B). Then, we tested whether DAP and P64 could regulate damaged protein loading and clearance. Indeed, DAP and P64 treatment resulted in the activation of protein synthesis (Figure 6D), and, consistent with this, the total protein content was also increased (Figure 6E). On the other hand, DAP and P64 treatment also resulted in a reduction in the proteasome activity (Figure 6F and Supplementary Figure 8C). Collectively, these results indicate that PDK1 is a potential target to enhance drug induced protein damage.

**Fig 6.**
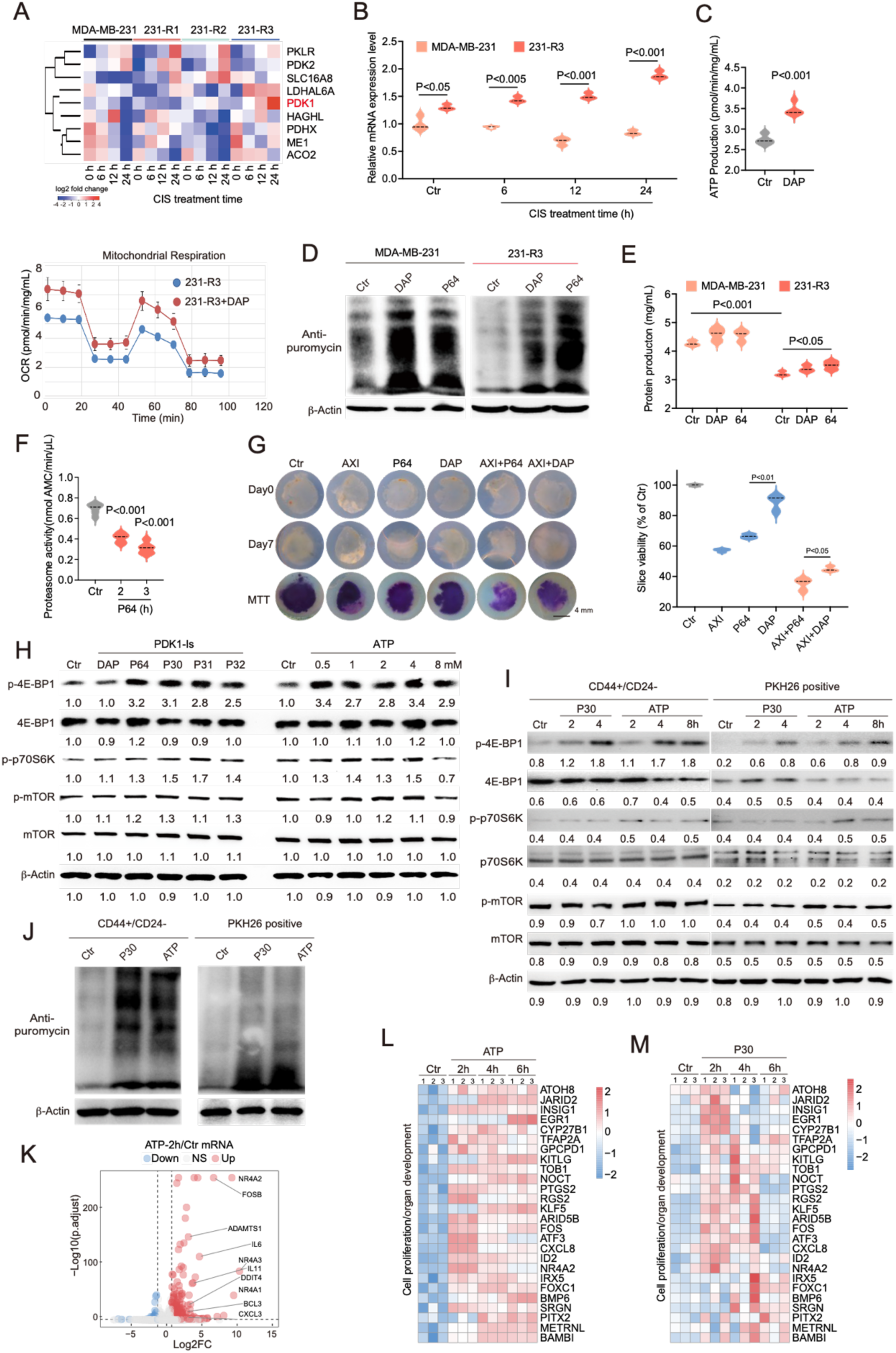
PDK1 inhibition forces ATP production that upregulates protein synthesis by enhancing p4E-BP1. (A) Nine up-regulated genes related to mitochondrial activity revealed by RNA-seq analysis of MDA-MB-231, 231-R1, 231-R2, and 231-R3 cells that were treated with *CIS*. (B) MDA-MB-231 and 231-R3 cells were treated with *CIS* for indicated time, and expression of PDK1 was detected by q-PCR analysis. (C) 231-R3 cells were treated with DAP, and mitochondrial OXPHOS and ATP production were measured by Seahorse XFp Cell Mito Stress Test assays. (D, E) MDA-MB-231 and 231-R3 cells were treated with DAP or P64, and protein synthesis was measured by Western blot (D) and total protein production was detected by BCA protein assay kit (E). (F) 231-R3 cells were treated with P64, and proteasome activity was measured by fluorescent AMC tagged peptide substrate. (G) 3D-TSCs were treated with *AXI* or combined with DAP or P64 treatment, and slice viability was measured by MTT assay. (H) MDA-MB-231 cells were treated with indicated PDK1-Is or ATP, and p-p70S6K at Threonine 389, p-4E-BP1 at serine 65, 4E-BP1, p-mTOR at serine 2448, and mTOR were detected by Western blot. (I, J) CD44+/CD24- cells and PKH26-positive cells that isolated from MDA-MB-231 were treated with P30 or ATP, and p-p70S6K at Threonine 389, p-4E-BP1 at serine 65, 4E-BP1, p-mTOR at serine 2448, and mTOR were detected by Western blot (I) and protein synthesis rate was measured by Western blot (J). (K, L) MDA-MB-231 cells were treated with ATP, and upregulated genes and down regulated genes were shown, p<0.05 (K), genes of cell proliferation and differentiation for 2-, 4-, and 6-hours were show by heatmap (L). (M) MDA-MB-231 cells were treated with P30, and genes of cell proliferation and differentiation for 2-, 4-, and 6-hours were show by heatmap. All values are presented as mean value (at least three replications) ± SD, and p value was calculated by comparison with Ctr group or indicated separately.

Next, we compared the efficacy of DAP and P64 in three-dimensional tumor slice culture (3D-TSC) drug sensitive test system (*42*), which is prepared from nude mice bearing the 231-R3 cells that are highly resistant to many anticancer drugs (*3*). The results indicated that P64 was more potently inhibited tumor growth, as indicated by the much weaker MTT staining of the 3D-TSCs (Figure 6G), which should be the result of increased specificity for PDK1 (*41*). Encouraged by these results, we further modified P64 and identified a few new P64 derivatives, P30, P31, and P32, with increased efficiency and specificity to PDK1 (*43*). Then, we compared combination drug regimens with P64, P30, P31, and P32 to identify the most promising PDK1-I. The results indicated that the modified drugs P30, P31, and P32 had enhanced efficiency compared to that of P64, with P30 more efficacious than the other drugs in 3D-TSCs and 231-R3 cells (Supplementary Figure 8D and E). We further tested these PDK1-Is in HCT and A549 cells, in which the 4 PDK1-Is tested in this study forced ATP production (Supplementary Figure 8F) and increased protein synthesis (Supplementary Figure 8G).

To investigate the mechanism by which PDK1-Is and ATP induce increased protein synthesis, we monitored the mTOR pathway, which plays a critical role in regulating protein synthesis (*44*). We found while all 4 new PDK1-Is (P30, P31, P32 and P64) did not cause an obvious change in the expression/phosphorylation of mTOR-p70S6K, they significantly increased the phosphorylation of initiation factor 4E-binding protein 1 (p4E-BP1) at serine 65 (Figure 6H), which dissociates it from eIF4E and activates cap-dependent translation (*45*). Similarly, increased 4E-BP1 phosphorylation was observed upon treatment with all tested concentrations of ATP (Figure 6H). These data indicate that PDK1-Is and ATP promote protein synthesis by enhancing 4E-BP1 phosphorylation. To investigate if this also occurs in CSCs, we sorted out CD44+/CD24- cells and PKH26-positive cells to enrich the CSCs from MDA-MB-231 cells and the data revealed that P30 and ATP treatment induced a time-dependent phosphorylation of 4E-BP1 in these cells while no obvious effect was observed on the phosphorylation of mTOR and p70S6K (Figure 6I). Consistently, the protein synthesis was increased by the treatment of P30 or ATP in the CSCs (Figure 6J)

Increased protein synthesis is the key feature of cell growth and fitness. We speculate that ATP could affect expression of genes involved cell proliferation. Indeed, inoculation of ATP to cells induced significant upregulation of gene expression in 2-, 4-, and 6-hours treatment, and very few genes were downregulated at mRNA levels (Figure 6K and Supplementary Figure 8H). The upregulated genes participate in the cell proliferation and differentiation, e.g. ossification, epithelial cell proliferation, and fat cell differentiation (Figure 6L and Supplementary Figure 8I). Consistently, ATP treatment also enhanced the protein level of genes involved in the nuclear division and chromosome segregation (Supplementary Figure 8J). These results imply that ATP could promote the cell proliferation. To validate this in the quiescent cells, MDA-MB-231 cells under serum starvation were treated with ATP in a time course manner, and we observed a significant increase of Ki67 staining in the early time point (Supplementary Figure 9A). Then, we treated quiescent MDA-MB-231-CG cells (obtained by serum starvation) with an increasing dose of ATP and found the entry of G1 cells into proliferative S phase (Supplementary Figure 9B).

Since PDK1 inhibition forced the ATP production, we tested these effects under P30 treatment. Similarly to ATP, P30 treatment enhanced expression of genes for cell proliferation and differentiation (Figure 6M). Consistently, in MDA-MB-231-CG cells, we observed that P30 treatment increased the fractions of proliferative cells (S or G2) and that the fraction of G1 cells eventually decreased (Supplementary Figure 9C). Thus, inhibition of PDK1 forces ATP production and promoted cells entry into the proliferative phase of the cell cycle, and both effects should enhance protein damage, as we demonstrated earlier. Consistently, combination treatment with DAP or P30 with 4 structurally different anticancer drugs resulted in increased protein damage (Supplementary Figure 9D). In summary, our results indicate that PDK1 is a potential target to reverse the low-ATP state that forces the CSCs and DRACs to proliferate and therefore render them more vulnerable to drug induced protein damage.

### Forced ATP production with proteasome inhibition to overcome MDR

We have previously shown that while proteasome-I treatment could reverse the resistance of the MDR cell line (231-R3) to many anticancer drugs, the 231-R3 line remains resistant to *temsirolimus* (*TLM*) or *atazanavir* (*ATV*) (*3*). Having identified PDK1 as a potential target to increase protein damage, we further formulated the “drugs + proteasome-I + PDK1-I” strategy to overcome the deeper level of MDR, and the data showed that this stubborn resistance to *TLM* and *ATV* was reserved by further combination with PDK1-I (Figure 7A and Supplementary Figure 10A). To further demonstrate this in vivo, 3-dimentional tumor slice culture (3D-TSC) (*46*) were prepared from nude mice bearing 231-R3 cells. The efficacy of the single drug was dramatically increased by proteasome-I treatment, which was further enhanced by the PDK1-I treatment (Figure 7B). Then, we validated whether targeting PDK1 with our combination treatment regimen could also regulate ATP production in vivo. We found that the ATP production was significantly increased by the P30 monotreatment of 3D-TSCs compared to the Ctr group, and similar results were also observed in the *CIS*+P30 group compared to the *CIS* group and in the *CIS*+*BTZ*+P30 group compared with the *CIS*+*BTZ* group (Figure 7C). This was accompanied by the increased protein synthesis in the P30 group compared with the Ctr group, *CIS*+P30 group compared with the *CIS* group, and *CIS*+*BTZ*+P30 group compared with the *CIS*+*BTZ* group (Figure 7D). Next, we tested proteasome activity. Since both P30 and *BTZ* can inhibit proteasome activity, we observed a significant decrease in proteasome activity in the *CIS*+P30, *CIS*+*BTZ*, and *CIS*+*BTZ*+P30 groups (Supplementary Figure 10B). Collectively, our data indicate that a treatment strategy consisting of “drugs + proteasome-I + PDK1-I” could overcome MDR by regulating ATP production and inducing protein damage.

**Fig 7.**
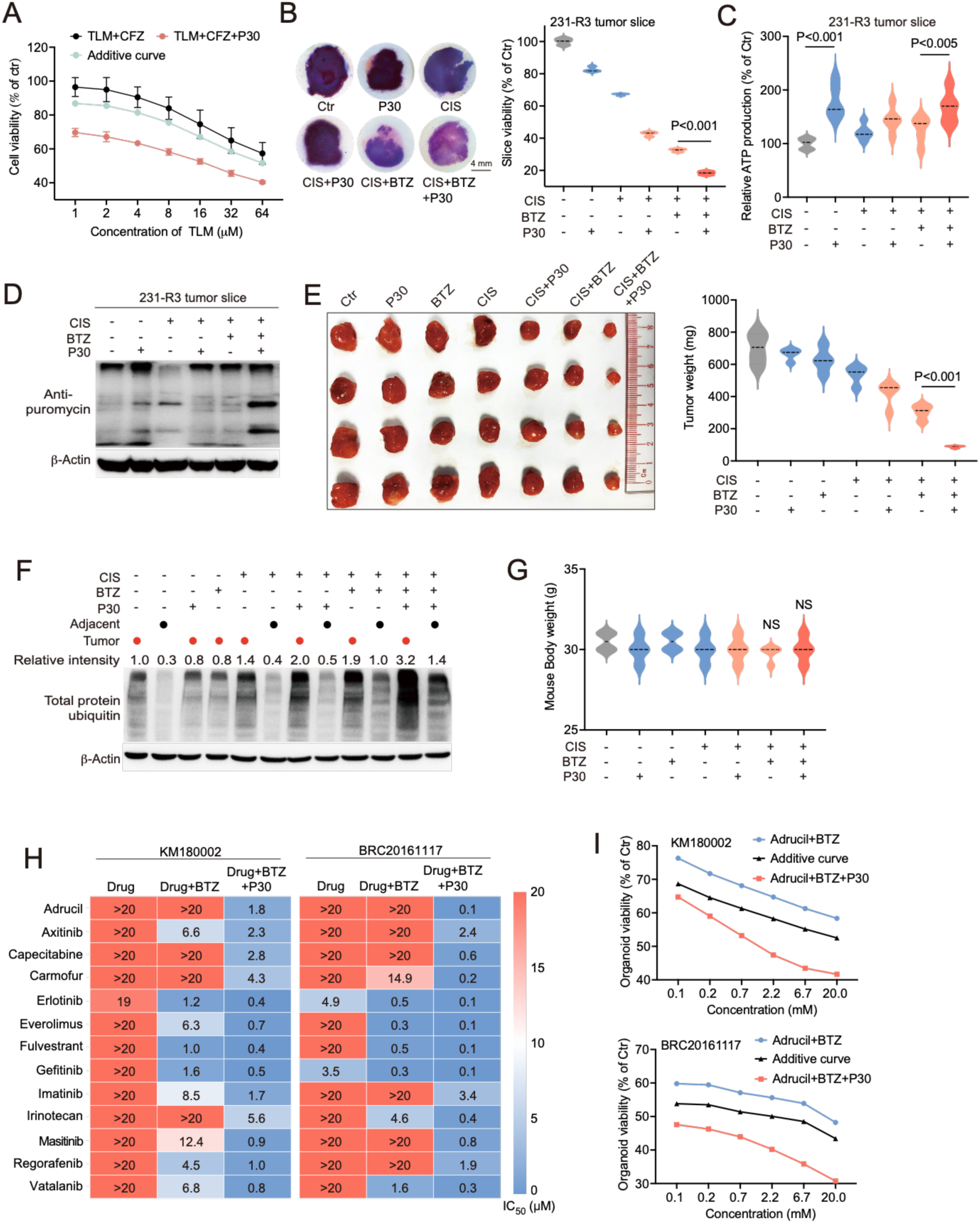
Forced ATP production with proteasome inhibition to overcome MDR. (A) Response of 231-R3 cells to *TLM* combined with non-toxic concentration of proteasome-I or *TLM* combined with non-toxic concentration of proteasome-I and PDK1-I. Cell viability was detected by alamar blue assay. Additive curves were calculated by Additive = E1+E2-E1*E2, where E1 is inhibition effect from *TLM*+*CFZ* and E2 is inhibition effect from P30 treatment. (B) The 3D-TSCs prepared from the 231-R3 xenograft tumors were treated with *CIS*, combine with PDK1-Is, or *BTZ*; each group is quintuplicate, and cell viability was measured by MTT assay. (C, D) The 3D-TSCs treated with *CIS* or combined with P30 or *BTZ* treatment, and ATP (C) levels and protein synthesis (D) were measured.(E to G) BALB/c mice bearing 4T1 allograft tumors were treated with vehicle, *CIS* (3 mg/kg), *BTZ* (1 mg/kg), or P30 (1 mg/kg) every 3 days for 3 times, and tumor photograph and tumor weight were recorded (E). Total protein ubiquitin for tumors and adjacent mammary gland were detected by Western blot (F). Body weight for mice were recorded (G). (H) 13 resistant drugs were tested in combination with *BTZ* or P30 treatment in KM180002 and BRC20161117 PDOs, and IC50 of 13 resistant drugs were shown for single drug treatment, double treatment, and triple treatment, respectively. (I) 1 most additive drug for P30 in KM180002 and BRC20161117 were demonstrated. All values are presented as mean value (at least three replications) ± SD, and p value was calculated by comparison with Ctr group or indicated separately.

Next, we validated our findings in BALB/c mice bearing 4T1 allograft tumors. We observed the significant inhibition of tumor growth and a reduction in tumor weight in the *CIS*+*BTZ* and *CIS*+*BTZ*+P30 groups, and these effects were associated with increased protein damage in these tumors (Figure 7E and Supplementary Figure 10C). To visualize the ATP content in situ, we conducted immunohistochemical staining with an antibody against ATP, and the data revealed a significant increase in ATP content in the *CIS*+P30 and *CIS*+*BTZ*+P30 tumors (Supplementary Figure 10C), which is consistent with the results obtained with an ATP detection kit (Supplementary Figure 10D). Cancer cells are characterized by sustained proliferation and frequent entry into the growth-and-division cycle, which is carefully controlled in normal tissues (*27*). This phenomenon places a heavy load on the translation machinery in cancer cells. Our earlier data indicated that anticancer drug-induced protein damage mainly targets ribosome-mediated protein synthesis, which might result in more severe damage to cancer cells and less severe damage to normal healthy tissues. Indeed, our treatment strategy induced a significant increase in protein damage and caused much less protein damage in adjacent normal tissue (Figure 7F), indicating minimal side effects. Consistently, the body weight of mice in the combination treatment group was similar to that of mice in the Ctr group (Figure 7G).

Patient-derived organoids (PDOs) has been shown to be reliable systems for cancer drug discovery and assessment (*47*). We have established culture conditions for PDOs from various cancers, including human breast and colon cancers (*48*). We previously showed the “drugs + proteasome-I” strategy for overcoming the resistance to 13 FDA approved drugs in 4 PDOs from colon cancer, while we noticed one PDO (KM180002) still resistant to such strategy (*3*). Then, we tested the “drugs + proteasome-I + PDK1-I” strategy in the KM180002 and the MDR of which is completely reversed (Figure 7H), and they showed the additive effect in the combination with PDK1-I treatment (Figure 7I). Similar results were observed in another refractory PDO (*3*) that was derived from a triple negative breast cancer at late stage (BRC20161117, Figure 7H, I), indicating the potential of “drugs + proteasome-I + PDK1-I” strategy for multitype of cancers.

## Discussion

MDR in cancer occurs very frequently and can be affected by many factors, including DNA damage repair, drug target mutation, drug metabolism and inactivation, cellular drug excretion, aberrant cell death regulation, and CSCs (*7, 8*). In this study, we found that vast majority of anticancer drugs damage proteins prior to reaching their canonic targets and induced protein damage response (PDR) that which is rarely recognized yet plays a critical role in MDR. We also demonstrated that the PDR contains two major steps, i.e. the damage recognition, which is mainly mediated by ubiquitin system and the damage clearance, which is mainly mediated by proteasome system as the treatment of proteasome inhibitors could result in the accumulation of protein damage, leading to the accelerated cell death.

We found that quiescent cells and slow proliferating cells are more resistant to protein damage and the associated lethality mainly because protein synthesis is relatively inactive in these cells compared to proliferative cells. We provide strong evidence that mitochondrial ATP production plays a key role in the regulation of both the protein synthesis rate and proteasome activity; therefore, the low-ATP status of these cells ensures low levels of protein synthesis and high levels of proteasome activity, both of which account for the drug resistance in these cells. Numerous studies have shown that CSCs are, in general, more resistant to chemotherapy than somatic cancer cells (*49–51*). Because dormancy is the key feature of CSCs (*21, 52–54*), our findings that CSCs and dormant cells exhibit lower levels of ATP production and protein synthesis and higher levels of proteasome activity provide a molecular basis for their drug resistance.

Acquired drug resistance commonly occurs during and after treatment through various mechanisms (*7*). To identify genes that may be responsible for this acquired drug resistance, we previously used RNAi libraries with increasing sizes to conduct screens. Our first RNAi library contained 56 genes, from which we identified ATP7A, which excretes cisplatin from cells, as a potential target that confers cisplatin resistance (*9*). We then screened an RNAi library containing 706 kinases and identified ATR, CHK1 and WEE1, which shut down DNA replication and attenuate cisplatin-induced lethality (*10*). While these screens confirmed that many factors may be involved in drug resistance, they were limited to identifying the most potent mechanisms underlying the generation of drug resistance because of the small number of genes screened. Thus, to identify the strongest molecular basis underlying cisplatin resistance, we finally applied he whole-genome RNAi screening together with tumor drug resistance evolution models and found that cisplatin resistance is closely related to the ability of cancer cells to remove damaged proteins (*3*). In proliferating cells, drug treatment induces extensive protein damage, resulting in strong cytotoxicity. On the other hand, because 99% of mitochondrial proteins are imported from the cytoplasm (*55*), the impairment of mitochondrial OXPHOS and shut down of ATP production due to these damaged proteins are unavoidable, which not only reduces new protein synthesis but also activates proteasome activity to reduce the loading of damaged proteins and enable cells to gradually acquire drug resistance.

Regarding the mechanism by which ATP stimulates protein synthesis, we found that both PDK1-Is and ATP increased p4E-BP1, which enhances cap-dependent translation and is over expressed in some types of human cancers (*45, 56*). 4E-BP1 is a well-known substrate of mTOR (*44*); however, we found that PDK1-I and ATP treatment did not cause an obvious change in mTOR-p70S6K, highlighting an mTOR-independent mechanism underlying the phosphorylation of 4E-BP1. Recent studies revealed that some other kinases, including GSK3β, p38MAPK, UVB, ERK, PIM, ATM, insulin, CDK1, LRRK2, and mTORC2, might phosphorylate 4E-BP1 at distinct sites under different conditions (*45*). Identifying the upstream kinases that phosphorylate 4E-BP1 and how they are affected by ATP is beyond the scope of this study; however, it might be an interesting topic for future investigation, as increased levels of p4E-BP1 have been detected in many different types of cancers.

Because a low-ATP state plays a key role in dictating drug resistance, we have developed a strategy to enhance mitochondrial respiration through the inhibition of PDK1 and enhancement of ATP production (*40*). Our data indicated that this could be effectively achieved by using DAP or several PDK1-Is that we recently developed (*43*). We further showed that enhanced mitochondrial ATP production activates dormant resistant cells and increases protein damage, with damaged proteins further accumulating upon blockade of damaged protein clearance with proteasome-Is to overcome MDR both in vitro and in vivo. Cancer cells, in general, divide faster than the cells in normal tissues and organs, and many cancers usually occur more frequently in older individuals, in which vast majority of noncancerous cells are less proliferative (*27, 57*). We also showed that our strategy is less toxic to healthy tissues due to their decreased degree of cell division and the presence of low baseline proteotoxic stress. Thus, targeting the damaged protein clearance pathway with our combination drug regimen serves as a promising and safe strategy to eliminate refractory cancer cells.

In summary, our data indicate that majority of anticancer drugs kill cells by damaging newly synthesized proteins and, at the same time, cancer cells initiate PDR to minimize the cytotoxicity as a mechanism to combat these drugs. The PDR includes protein damage recognition and clearance, and it also downregulates mitochondrial activity/ATP production, which not only reduces protein synthesis in these cells by enhancing p4E-BP1 but also upregulates proteasome activity, enabling them to acquire MDR (Supplementary Figure 11). On the other hand, quiescent cells, including CSCs, maintain low-ATP status; therefore, due to their lower levels of protein synthesis and higher levels of proteasome activity, these cells are intrinsically resistant to the cytotoxicity of anticancer drugs. We also showed that drug resistance could be reversed through a forced increase in mitochondrial ATP production, which enhanced protein synthesis and disrupted cell dormancy, rendering cancer cells more vulnerable to the cytotoxicity of anticancer drugs. Because MDR is a major problem and kills the majority of cancer patients (*6–8*), our study may provide a promising strategy to determine the drug resistance status of cancer patients before, during and after treatment by measuring their ATP levels and/or proteasome activity; furthermore, proper therapeutic options may be designed accordingly.

## Methods

### Mouse strains

All mouse experiments were performed under the ethical guidelines of the University of Macau (animal protocol number: UMARE-AMEND-219). Mice were housed in a specific-pathogen-free (SPF) facility at 23–25 °C on a 12-h light/dark cycle. The mice were fed a standard rodent diet (PicoLab Rodent 5053 Laboratory Diet). BALB/c mice and nude mice (5-8 weeks) were used in this study.

### Allograft/xenograft studies

MDA-MB-231 and 231-R3 multidrug-resistant cells (derived from MDA-MB-231 cells) were cultured in DMEM with 10% fetal bovine serum. The cells were trypsinized and resuspended in PBS. Then, the cells were mixed with Matrigel (Corning, 354234) at a 1:1 (v/v) ratio, and 1.0 × 10^6^ cells were injected subcutaneously into the flanks of nude mice. 4T1 cells were cultured in RPMI 1640 medium with 10% fetal bovine serum. The cells were prepared as above, and 1.0 × 10^6^ cells were injected into the mammary fat pads.

Tumor volume was calculated as V = (W^2^ × L)/2. The mice began receiving drug treatment when the tumor volume reached ∼200 mm^3^. Cisplatin (3 mg/kg), BTZ (1 mg/kg), CFZ (1 mg/kg), P30 (1 mg/kg), palbociclib (25 mg/kg), artemether (10 mg/kg), and lapatinib (25 mg/kg) were administered by intraperitoneal injection.

### Cell culture

T47D, MCF7, MDA-MB-231, HepG2, A549, HCT, TM91, JIMT-1, HCC1954, and SKBR3 cells were cultured in DMEM supplemented with 10% fetal bovine serum, glutamine (Thermo), and pen/strep. The cisplatin-resistant cell lines 231-R1, 231-R2, and 231-R3 were established as previously described (*3*).

### 3D-TSCs

3D-TSCs were prepared from allograft and xenograft tumors. Thick tissue slices (200 μm) were prepared under sterile conditions using a Leica VT1200 S vibratome (Leica Biosystems Nussloch GmbH, Germany) within 2 h after surgery. An embedding collagen mixture containing rat collagen I, 10X Ham’s F-12 medium, and sterile reconstitution buffer (2.2 g NaHCO_3_ in 100 ml of 0.05 N NaOH and 200 mM HEPES) at a ratio 8:1:1 was prepared. The thick tissue slices were embedded in 100 μL of the collagen mixture on the bottom and 50 μL of the collagen mixture on the top. The slices were then placed on 0.4-mm pore-size membrane culture inserts in 24-well plates and cultured in F medium, which contained DMEM/F12 (Thermo), 5% FBS, 10 ng/ml EGF (Thermo, 13247-051), 0.5 mg/ml hydrocortisone (Sigma, H0888), 20 ng/ml cholera toxin (Sigma, C-3012), 5 μg/ml insulin (Sigma, 350-020), 300 U/ml collagenase III (Worthington, S4M7602S) and hyaluronidase (Sigma, H3506). Drug treatment was performed after overnight culture, and images were taken with a Leica M165FC fluorescence stereomicroscope. For MTT staining, the dye MTT (3-[4,5-dimethylthiazol-2-yl]-2,5-diphenyltetrazolium bromide; thiazolyl blue, Sigma[Aldrich) was added (final concentration: 0.5 mg/ml) to tissue slices and incubated for 4 h. The formazan crystals were solubilized with DMSO, and the absorbance at 570 nm was measured using a plate spectrophotometer.

### Chemicals and anticancer drug library

The anticancer drug library used to screen for protein damage and DNA damage and chemicals used in this study are summarized in Supplementary Table. Other chemicals included puromycin (InvivoGen, ant-pr-1), CHX (biorbyt, orb340214a), salubrinal (Selleckchem, S2923-5mg), NAC (Beyotime, S0077), thymidine (Sigma[Aldrich, T9250), L-ascorbic acid (Beyotime, ST1434-25g), CFZ (MedChemExpress, HY-10455/50mg), BTZ (MedChemExpress, HY-10227/100mg), IXZ (MedChemExpress, HY-10453/25mg), Doxycycline (Sigma–Aldrich, D9891), adenosine 5′-triphosphate disodium salt hydrate (Sigma[Aldrich, A2383-10G), oligomycin (Sigma[Aldrich, 75351), antimycin A (Sigma[Aldrich, A8674), 2,2-dichloroacetophenone (Sigma[Aldrich, D54850), biotin (MedChemExpress, HY-B0511), cisplatin-biotin (Xi’an ruixi Biological Technology Co.,Ltd, R-HGF148), and streptavidin agarose (Sigma-Aldrich, 85881).

### Cell viability analysis

Cell viability was determined by the alamar blue assay. Briefly, cultured cells were plated in a 96-well plate at 5,000 cells per well and cultured overnight. The cells were treated with the indicated drugs for 48 h. Then, the cells were incubated with medium containing 0.02% alamar blue for 2 h, and cell viability was determined by measuring the absorbance at 590 nm under 560-nm excitation.

### TCGA/TARGET database analysis

The present study leveraged a pan-cancer dataset from the TCGA and TARGET databases. Upper-quartile normalized FPKM RNA-sequencing data were retrieved from the UCSC XENA resource (*58*), while clinical data were acquired through the R package UCSCXenaTools. Subsequently, proteasome activity score was quantified for each sample using the Gene Set Variation Analysis (GSVA) function implemented in the gsva R package (*59*).

### Serum starvation

MDA-MB-231, 4T1, and HepG2 cells were cultured in 0.01% fetal bovine serum for 3 days to enter the quiescent G0 state. MCF-7, HCT, and A549 cells were cultured in 0.1% fetal bovine serum for 3 days to enter the quiescent G0 state.

### Quantitative real-time PCR

PDK1 expression in MDA-MB-231 and 231-R3 cells was confirmed using real-time PCR. The primer sequences used were as follows: forward CTGTGATACGGATCAGAAACCG and reverse TCCACCAAACAATAAAGAGTGCT.

### Western blot and immunoprecipitation

Cultured cells were homogenized in RIPA buffer supplemented with phosphatase and protease inhibitor cocktails (Sigma). The protein concentration was detected by a BCA protein assay kit, and samples containing equivalent protein concentrations were resolved by SDS/PAGE (10% gels). The proteins were then transferred to PVDF membranes (Bio-Rad), which were blocked with 5% BSA in TBS-T buffer for 1 h at room temperature and incubated with primary antibody overnight at 4 °C. The PVDF membranes were incubated with the corresponding secondary antibody (Cell Signaling Technology, CST), and the protein bands were visualized with horseradish peroxidase substrate (Millipore) under a ChemiDoc Touch Imaging System (Bio-Rad). Primary antibodies against the following were used in this study: ubiquitin (1:1000, CST, 3936S), γH_2_AX (1:1000, CST, 9718S), β-actin (1:1000, CST, 3700S), HA (1:2000, Sigma–Aldrich, H6908), Myc (1:200, Santa Cruz Biotechnology, sc-40), FLAG (1:2000, Sigma–Aldrich, F1804), cleaved caspase-3 (1:1000, CST, 9664S), anti-puromycin (1:1000, Merck, MABE343), GAPDH (1:1000, CST, 5174S), GFP (1:200, Santa Cruz Biotechnology, sc-9996), luciferase (1:200, Santa Cruz Biotechnology, sc-57603), p4E-BP1 (1:1000, CST, 9451), 4E-BP1 (1:1000, CST, 9452), p70S6K (1:200, Merck Millipore, 06-926), p-p70S6K (1:1000, CST, 9234), pAKT (1:1000, CST, 9271), mTOR (1:1000, CST, 2983), and p-mTOR (1:1000, CST, 5536).

For immunoprecipitation, cultured cells were homogenized in IP cell lysis buffer (Beyotime) and incubated with primary antibody and Protein A/G PLUS-Agarose (Santa Cruz Biotechnology, sc-2003) overnight at 4 °C. Proteins of interest were then pulled down by centrifugation (Eppendorf) and detected by Western blotting.

### Lentiviral short hairpin RNA (shRNA) transfection

shRNAs were cloned and inserted into the lentiviral vector pLKO.1 puro (Addgene Plasmid 10878) according to the manufacturer’s instructions. After transfection, positive cells were selected by puromycin (Thermo Fisher); the knockdown effect of the shRNAs was reported previously (*3*). The following sequences were used: NDUFS1, forward TAACCTTTGTGACGAACATAA, reverse TTATGTTCGTCACAAAGGTTA; NDUFA2, forward CGTGAGATTCGCATCCACTTA, reverse TAAGTGGATGCGAATCTCACG.

### Protein aggregation detection assay

The PROTEOSTAT aggresome detection kit (Enzo Life Sciences, ENZ-51035-K100) was used to detect mis-folded or aggregated proteins in cells treated with indicated anticancer drugs. The PROTEOSTAT staining was performed according to the manufacturer’s instructions. Briefly, MDA-MB-231 cells were treated with indicated drugs, fixed, and permeabilized. The PROTEOSTAT was observed by 488 excitation and 600-620 emission on the Carl Zeiss LSM880 Confocal.

### Protein carbonyl detection

Cultured cells were lysed, and proteins were extracted and processed using a Protein Carbonyl Content Assay Kit (Abcam, ab178020) according to the manufacturer’s instructions. Briefly, the carbonyl groups in the protein side chains were derivatized to 2,4-dinitrophenylhydrazone (DNP-hydrazone) by reaction with 2,4-dinitrophenylhydrazine (DNPH), and DNP-hydrazone was detected by Western blotting with primary anti-DNP antibody.

### Proteasome activity detection

Cultured cells, 3D-TSCs, or mouse tumor tissues were homogenized in 0.5% NP40, and proteasome activity was measured by a Proteasome Activity Assay Kit (Abcam, ab107921) according to the manufacturer’s protocol. Briefly, cell lysates were incubated with Suc-Leu-Leu-Val-Tyr-7-amino-4-methylcoumarin (Succ-LLVY-AMC), and the free AMC fluorescence was measured at 350/440 nm. The increase in AMC fluorescence (ΔRFU) was detected as proteasomal activity as follows: ΔRFU = (RFU2 – iRFU2) – (RFU1 – iRFU1), where RFUn is the total proteolytic activation at time n and iRFUn is the non-proteasomal proteolytic activity at time n.

### Bulk RNA sequence

MDA-MB-231 cells were treated with 1 mM ATP or 2 μM P30 for 2-, 4-, 6 hours, and harvested for RNA sequence. After accessing the raw expression matrixes of each group, genes with low expression were filtered out. We retained genes with an expression value greater than 1 count per million (cpm) in at least 50% of the samples from the expression matrix. Deseq2 was employed to build the statistical model, utilizing shrinkage estimation to improve the stability of the fold changes and dispersions (*60*). Differential expressed genes (DEGS) were calculated based on treated groups and control groups. The threshold for DEGs was set as an adjusted P (BH method) value < 0.01 and an absolute value of log_2_ fold_change > 1.

For functional enrichment analysis, the R package clusterProfiler (*61*) was utilized according to the different pathway knowledges like Gene Ontology (Go) (*62*).

### Proteomics

MDA-MB-231 cells were treated with 1 mM ATP for 3 and 9 hours, proteins were harvested and submitted to LC-MS/MS analyze. MDA-MB-231 cells were treated 50 μM biotin, 50 μM cisplatin-biotin, 50 μM cisplatin-biotin+20 nM BTZ, or 50 μM cisplatin-biotin+50 µg/ml CHX for 3 hours, and cells were lysed. Biotin was pull down by the streptavidin immobilized on the agarose beads. The binding proteins were analyzed by LC-MS/MS. The protein peaks data was utilized to represent the final expression levels of a specific protein across various samples, resulting in a 5106 x 9 and 3751 x 12 protein-expression matrix for ATP treatment and biotin treatment, respectively. Missing values were imputed as one-tenth of the minimum value across our proteome dataset. The Student’s t-test, as implemented in R software, was used to examine proteins differentially expressed between the following groups: ATP_3 h vs. Ctr, ATP_9 h vs. Ctr, Bio-CIS vs. Bio, Bio-CIS+CHX vs. Bio, and Bio-CIS+BTZ vs. Bio. Proteins were considered upregulated if they were differentially expressed compared to the Ctr or Bio group (fold change (expressed as log2(ratio of average protein abundance)) > 1, p < 0.05).

### Immunohistochemical staining

Routine haematoxylin staining of 6-μm sections of formalin-fixed, paraffin-embedded (FFPE) tissues was performed. IHC staining for cleaved caspase-3 (1:500, CST, 9664S), ubiquitin (1:100, Thermo Fisher, 14-6078-95), Ki67 (1:200, Abcam, ab16667), and ATP (1:76, Cloud-Clone, PAA349Ge01) was performed by a Histostain-Plus IHC Kit (Thermo, 859043, 1954379A) after antigen retrieval (pH 6.0), quenching of endogenous peroxidase activity and overnight incubation with the primary antibody.

### ATP detection

For cellular total ATP detection, ATP levels were measured using the Cell Titer-Glo Luminescent Cell Viability Assay (Beyotime, C0065M) according to the manufacturer’s instructions. Briefly, cultured cells were lysed to release ATP, and luciferase enzymes catalyzed the oxidation of luciferin into oxyluciferin in the presence of ATP, generating a luminescent signal proportional to the amount of ATP.

Live-cell ATP detection was performed by the Cell Meter™ Live Cell ATP Assay Kit (AAT Bioquest, 23015) according to the manufacturer’s protocol. ATP Red™ is a cell-permeable red fluorescent probe used to image and detect ATP in live cells, and the ATP Red™ signal was observed by an LSM880 confocal microscope (Carl Zeiss).

### Detection of mitochondrial OXPHOS and ATP production

To detect mitochondrial OXPHOS, the real-time OCR was measured on a Seahorse XF24 Analyzer (Agilent Technologies, USA). Cells were plated in 150 μL of complete DMEM at 1 × 10^5^ cells/well on 24-well Seahorse plates (Agilent Technologies, USA) 24 h prior to the assays. The medium was removed and replaced with glutamine- and pyruvate-free DMEM (D5030 pH 7.4). Mitochondrial OXPHOS and ATP production were calculated by XF24 Analyzer software (Agilent Technologies, USA) after 1.5 h of incubation in DMEM. To detect the effects of the drugs on mitochondrial OXPHOS and ATP production, cells were treated with the indicated drugs before measurement on a Seahorse XF24 Analyzer.

### Fluorescence-activated cell sorting

MDA-MB-231 and 231-R3 cells at a concentration of 1 × 10^6^ cells per 100 μL of FACS buffer (Hanks’ balanced salt solution with 0.5% BSA and 2 mm EDTA) were stained. Antibodies against human CD24 (BioLegend, 311106) and CD44 (BioLegend, 338806) were used. Antibody incubation was performed at 4 °C for 30 min. PKH26 (Sigma Aldrich, MINI26) staining was conducted according to the manufacturer’s protocol. Briefly, MDA-MB-231 and 231-R3 cells at 1 × 10^7^ cells/mL were incubated with 2 μM PKH26 for 5 min. After centrifugation and removal of the supernatant, the cells were cultured in DMEM with 10% foetal bovine serum for 2 weeks. Then, antibody- or PKH26-labelled MDA-MB-231 and 231-R3 cells were washed with PBS twice before sorting by using a FACSAria III Cell Sorter (BD Biosciences). CD44+/CD24-, CD44-/CD24-, PKH26-negative, and PKH26-positive cells were gated and sorted for the detection of ATP and protein synthesis.

### Figure construction

Graphs were created in Prism (v.9) and assembled into figures in Adobe Illustrator 2022. All experiments were conducted at least three times independently. Microsoft Excel and Prism (v.9) were used for statistical calculations, and t test analysis was used to determine the significance of the difference between different sets of data.

## Data availability

The datasets used and/or analyzed in the current study are available in Supplementary Table and the source data, and further information is available from the corresponding author upon reasonable request.

## Acknowledgements

We would like to thank the members of the C.X.D. and X.L.X. laboratories for helpful advice and discussions. We are grateful for the support of the Animal Research Core of the Faculty of Health Sciences for providing the animal housing.

---

**Supplementary Fig 1.**
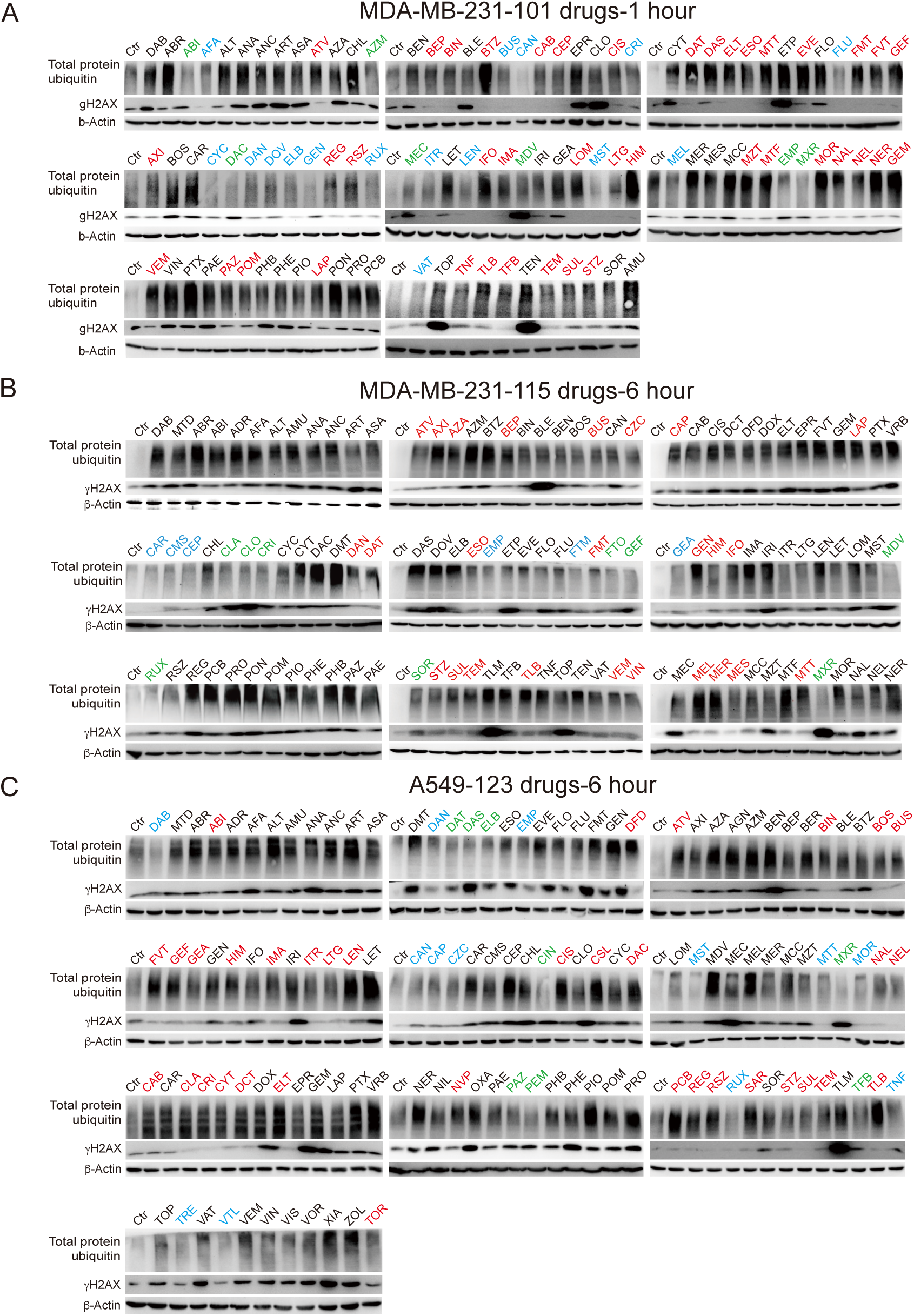
The vast majority of anticancer drugs cause protein damage prior to affect their canonic targets. (A, B) MDA-MB-231 cells were treated with 101 drugs for 1 hour (A) and 115 drugs for 6 hours (B), and protein total ubiquitin and γH_2_AX were detected by Western blot. Drugs only induce protein damage (red), drugs only induce DNA damage (green), and drugs induce both protein damage and DNA damage (black), and drugs do not damage protein and DNA (blue). (C) A549 cells were treated with 123 drugs for 6 hours, and protein total ubiquitin and γH_2_AX were detected by Western blot.

**Supplementary Fig 2.**
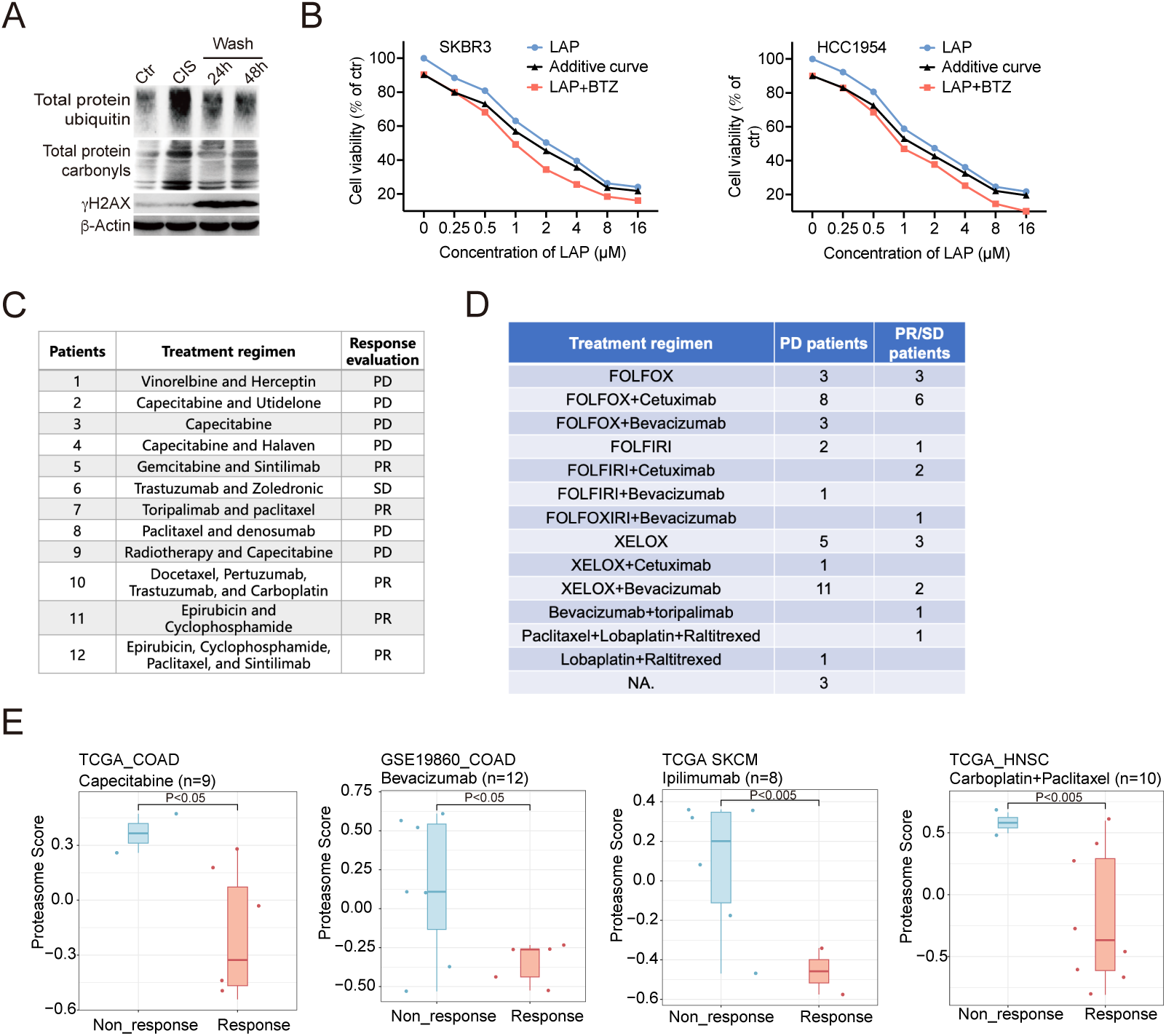
The vast majority of anticancer drugs cause protein damage prior to affect their canonic targets. (A) MDA-MB-231 cells were treated with *CIS* for 6 hours and washed away, and total protein ubiquitin, protein carbonylation, and γH_2_AX were detected by Western blot at 24 hours and 48 hours. (B) Two breast cancer cell lines were treated with *LAP* alone or combined with *BTZ* treatment, and cell viability was detected by alamar blue assay. Additive curves were calculated by Additive = E1+E2-E1*E2, where E1 is inhibition effect from *LAP* and E2 is inhibition effect from *BTZ* treatment. (C, D) The drug treatment regimen for 12 breast cancer patients (C) and 58 colon cancer patients (D), the drug response were evaluated by the changes of tumor burden according to the RECIST guideline (version 1.1). (E) Patients from TCGA and GES databases, the drug treatment response and the proteasome activity score. All values are presented as mean value (at least three replications) ± SD, and p value was calculated by comparison with Ctr group or indicated separately.

**Supplementary Fig 3.**
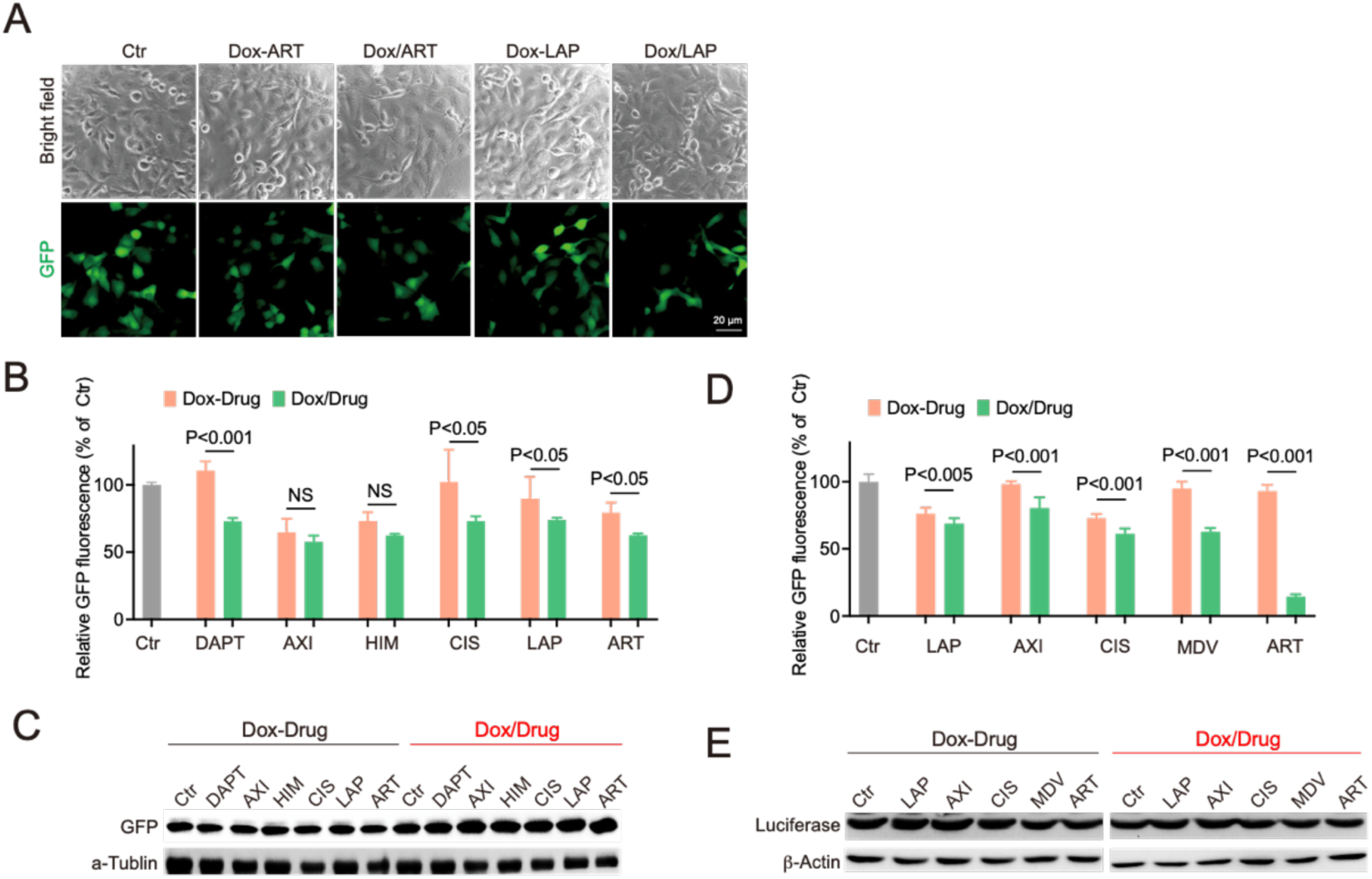
Anticancer drugs damage neosynthesized proteins and impair their functions. (A-C) MDA-MB-231-GFP cells were treated with indicated drugs, and GFP fluorescence was recorded (A). GFP fluorescence were measured by microplate reader (B), and expression of GFP was detected by Western blot (C). (D, E) MDA-MB-231-luciferase cells were treated with indicated drugs, and luciferase activity were measured by microplate reader (D), and expression of luciferase was detected by Western blot (E). All values are presented as mean value (at least three replications) ± SD, and p value was calculated by comparison with Ctr group or indicated separately.

**Supplementary Fig 4.**
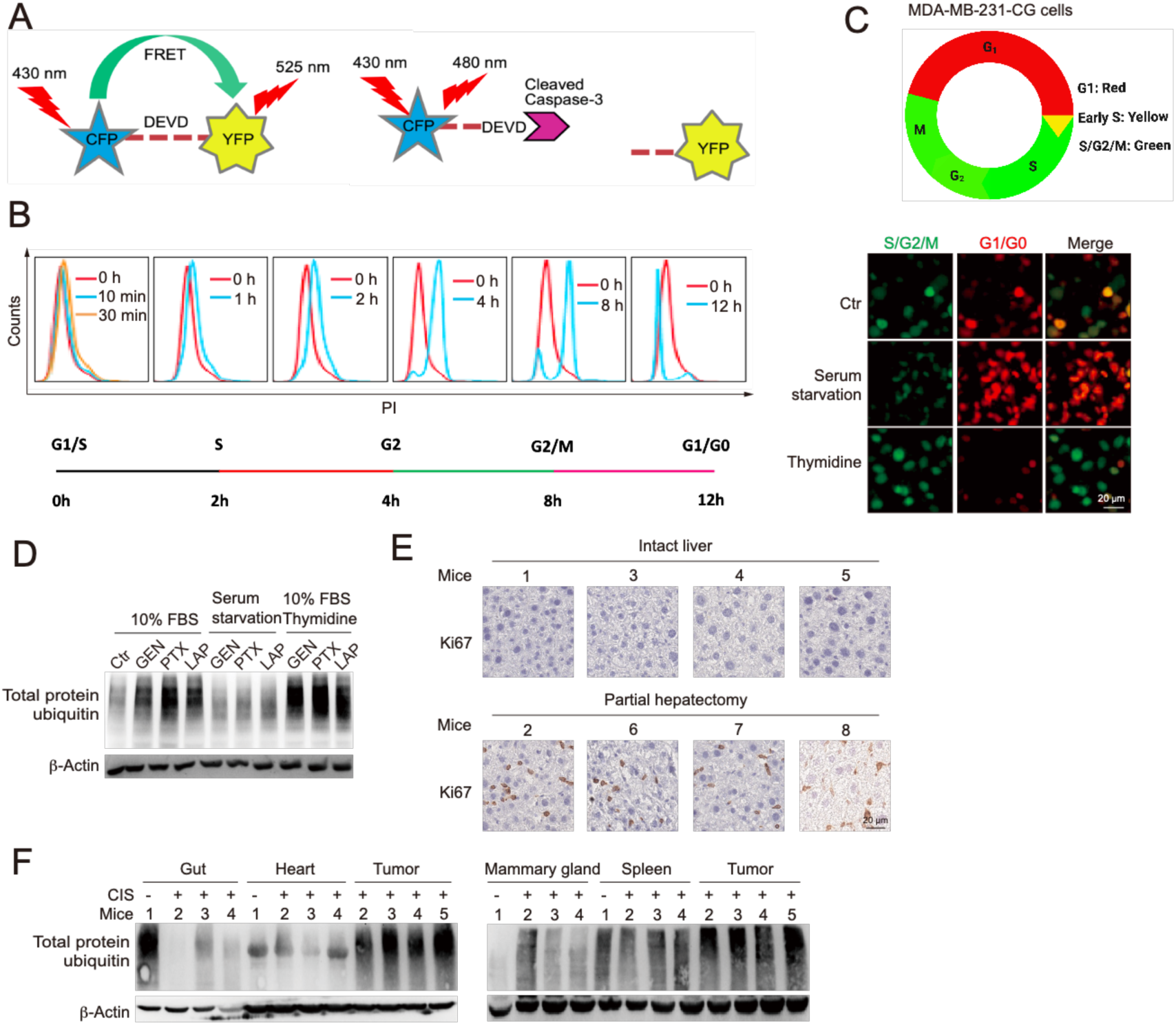
Protein damage occurs at low levels in quiescent cells but high levels in proliferating cells. (A) Principle of caspase-3 reporter Sensor C3. (B) MDA-MB-231 cells were synchronized by double thymidine treatment, and release at 10 min, 30 min, 1 hour, 2 hours, 4 hours, 8 hours, and 12 hours, and DNA content was measured by flow cytometry. (C and D) MDA-MB-231-CG cells were labeled with mKO2-Cdt1 (expressed at G1, red), mAG-Geminin (expressed at S/G2/M, green), and early S phase express both mKO2-Cdt1 and mAG-Geminin (yellow). MDA-MB-231-CG cells were under quiescent state (FBS starvation) or thymidine treatment, and cell cycle was recorded (C) and protein ubiquitin was monitored by Western blot (D). (E) Eight 5-month-old mice with or without partial hepatectomy were treated with *CIS* (6 mg/kg) for 24 hours, and Ki67 expression in the liver was detected by immunohistochemistry. (F) BALB/c mice carrying 4T1 allograft tumors were treated with *CIS* (6 mg/kg) for 24h, and total protein ubiquitin for tumors, adjacent mammary gland, and several organs were detected by Western blot.

**Supplementary Fig 5.**
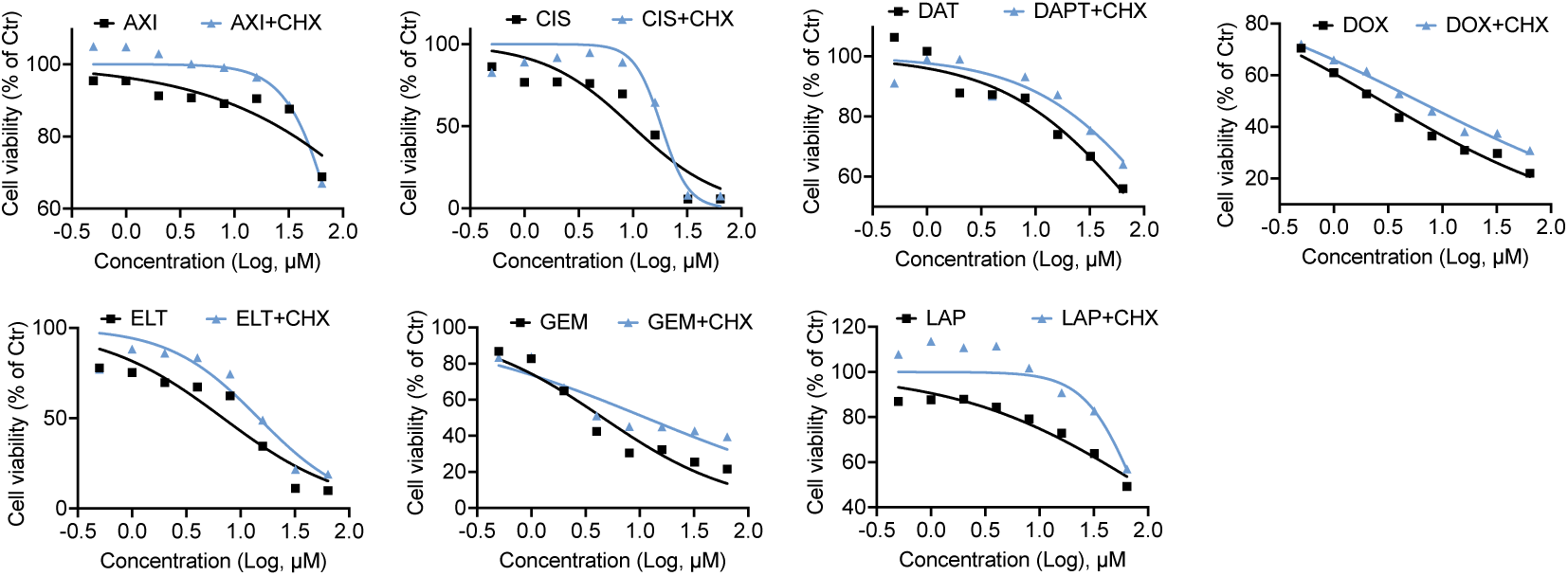
Effect of CHX on drug sensitivity. MDA-MB-231 cells were treated with indicated drugs or combined with CHX treatment, and cell viability was detected by alamar blue assay.

**Supplementary Fig 6.**
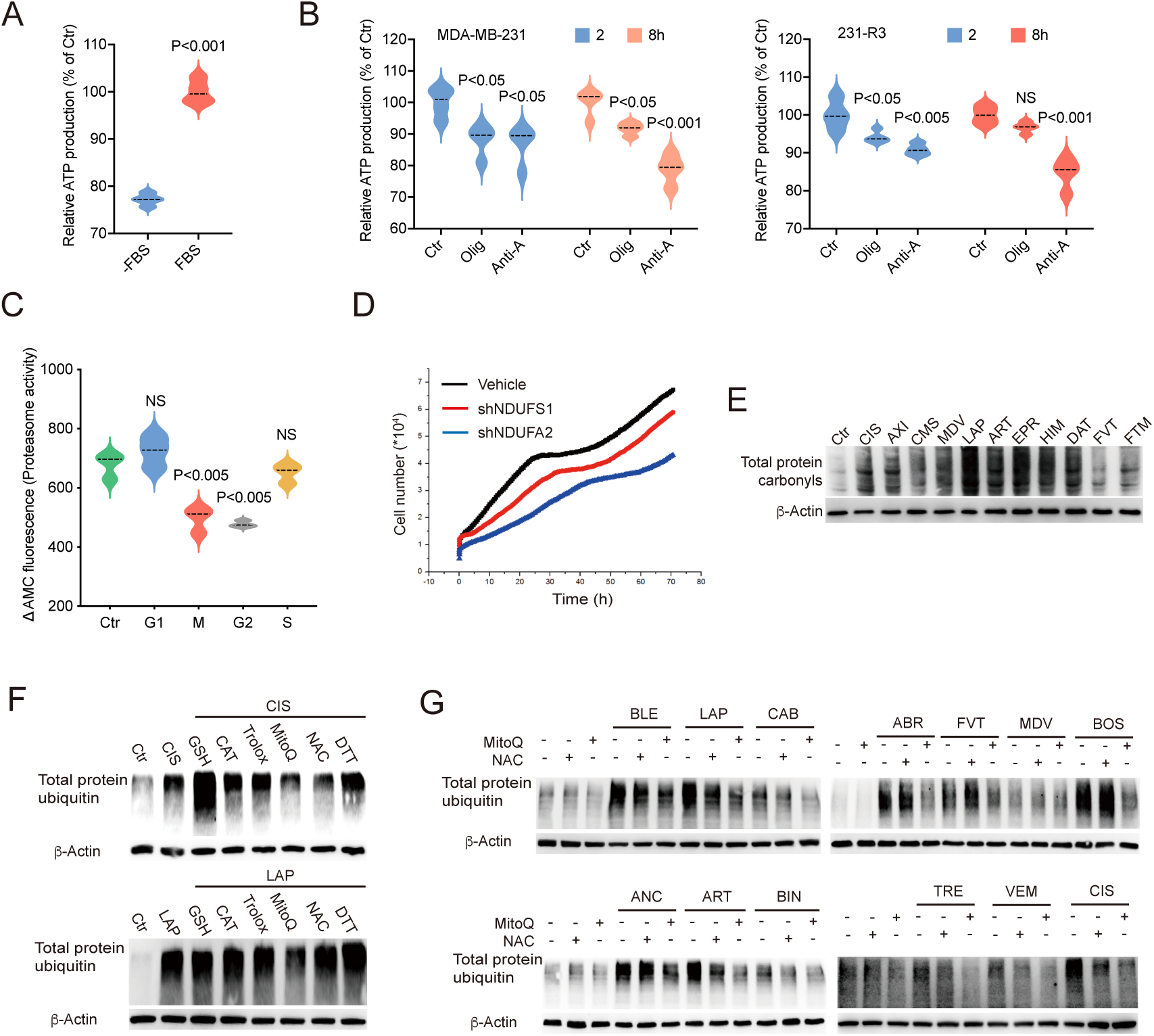
Maintenance of a low-ATP status is a common strategy by which CSCs and drug-resistant acquired cells (DRACs) acquire MDR. (A) MDA-MB-231 quiescent cells and cells under 10% FBS culture were tested for ATP production by ATP detection kit. (B) MDA-MB-231 and 231-R3 cells were treated with oligomycin or antimycin A, and ATP quantity is detected by ATP detection kit. (C) The proteasome activity for cells at S phase, G2 phase, M phase, and G1 phase were measured by fluorescent AMC tagged peptide substrate. (D) cell proliferation rate of MDA-MB-231 cells after knocking down of mitochondrial respiratory chain genes. (E) MDA-MB-231 cells were treated with indicated anticancer drugs for 6 hour, and total protein carbonylation was detected by Western blot. (F) MDA-MB-231 cells were treated with indicated drugs or combined with GSH, DTT, Trolox, NAC, ROS catalase, or MitoQ, protein total ubiquitination was detected by western blot. (G) MDA-MB-231 cells were treated with indicated drugs or combined with NAC or MitoQ, protein total ubiquitination was detected by western blot. All values are presented as mean value (at least three replications) ± SD, and p value was calculated by comparison with Ctr group or indicated separately.

**Supplementary Fig 7.**
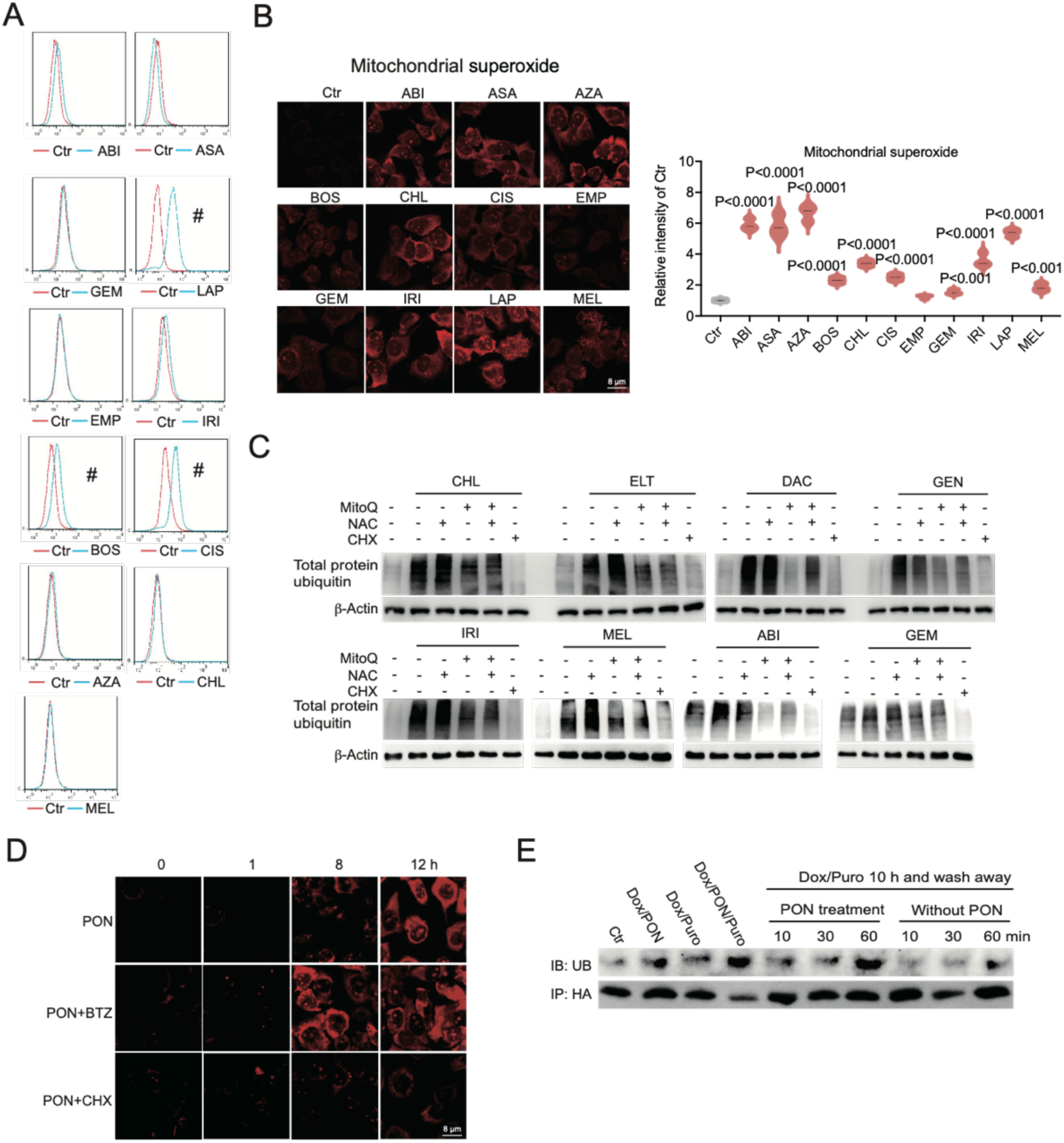
Maintenance of a low-ATP status is a common strategy by which CSCs and drug-resistant acquired cells (DRACs) acquire MDR. (A, B) MDA-MB-231 cells were tested with 11 drugs, and cytosol ROS accumulation was detected by DCFDA staining (A), and mitochondrial superoxide were detected by MitoSOX™ Red reagent (B). (C) MDA-MB-231 cells were treated with indicated drugs or combined with NAC, MitoQ, or CHX, and protein total ubiquitination was detected by western blot. (D) MDA-MB-231 cells were treated with *PON* alone or combined with *BTZ* or CHX treatment, and mitochondrial superoxide were detected by MitoSOX™ Red reagent. (E) MDA-MB-231-GFP cells were treated with dox and puromycin for 10 h to generate the nascent peptides, followed by incubation with or without *PON* treatment for indicated time, and GFP was pulled down for detection of ubiquitination.

**Supplementary Fig 8.**
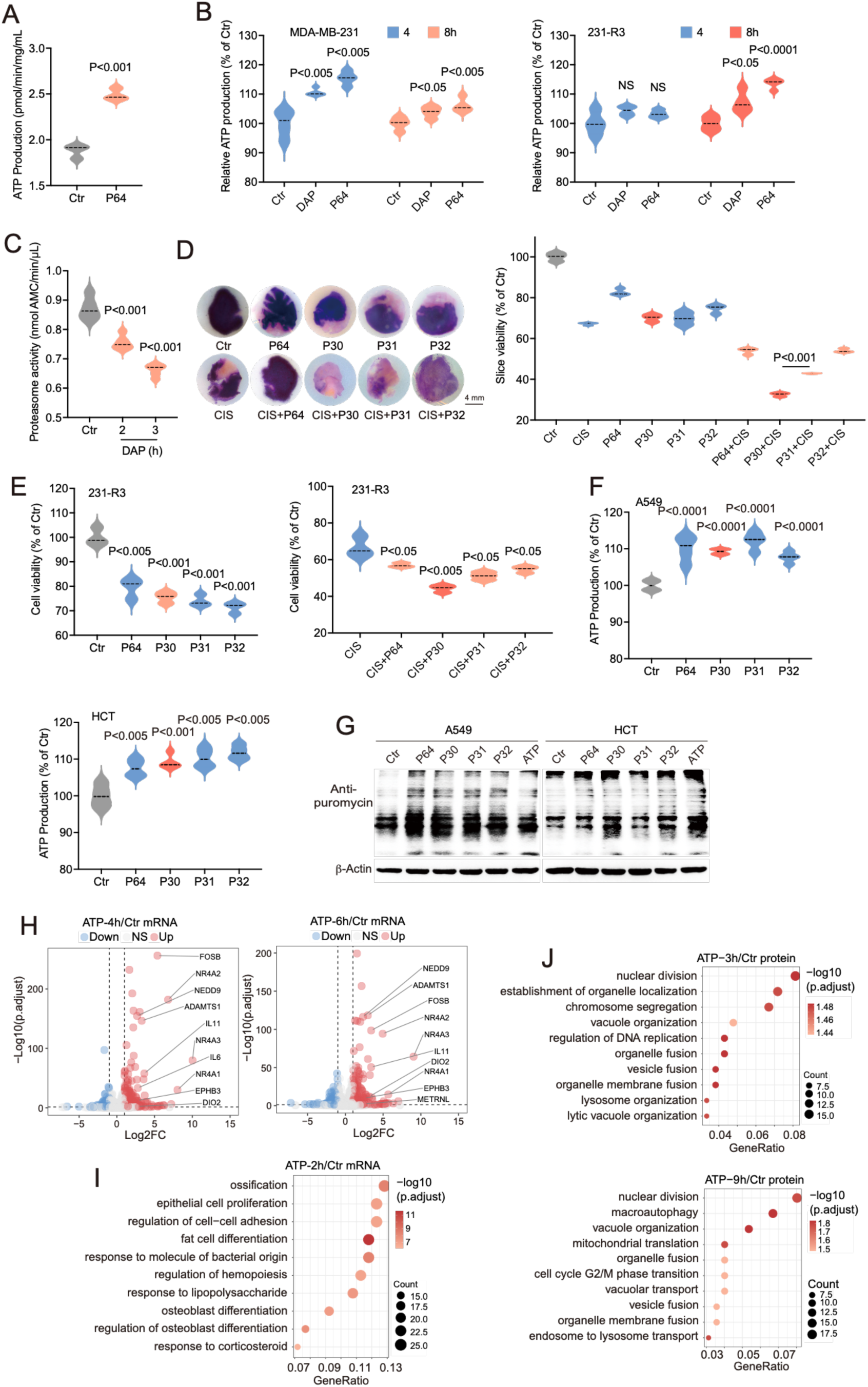
PDK1 inhibition forces ATP production that upregulates protein synthesis by enhancing p4E-BP1. (A) 231-R3 cells were treated with P64, and mitochondrial ATP production were measured by Seahorse XFp Cell Mito Stress Test assays. (B) MDA-MB-231 and 231-R3 cells were treated with DAP or P64, and ATP concentration was detected by ATP assay kit. (C) 231-R3 cells were treated with DAP, and proteasome activity was measured by fluorescent AMC tagged peptide substrate. (D) The 3D-TSCs prepared from the 231-R3 xenograft tumors were treated with *CIS*, combine with PDK1-Is; each group is quintuplicate, and cell viability was measured by MTT assay. (E) 231-R3 cells were treated with *CIS*, or combined with 4 different PDK1-Is, and cell viability is detected by alamar blue assay. (F, G) A549 and HCT cells were treated with indicated PDK1-Is for 8 hours, and ATP concentration was detected by ATP assay kit (F). Protein synthesis was detected by Western blot (G). (H, I) MDA-MB-231 were treated with ATP for indicated time, and upregulated genes, and down regulated genes were shown, p<0.05 (H), up regulated genes were analyzed by GO term enrichment (I). (J) MDA-MB-231 were treated with ATP for indicated time, and protein level were detected by mass spectrometry, and up regulated genes were analyzed by GO term enrichment. All values are presented as mean value (at least three replications) ± SD, and p value was calculated by comparison with Ctr group or indicated separately.

**Supplementary Fig 9.**
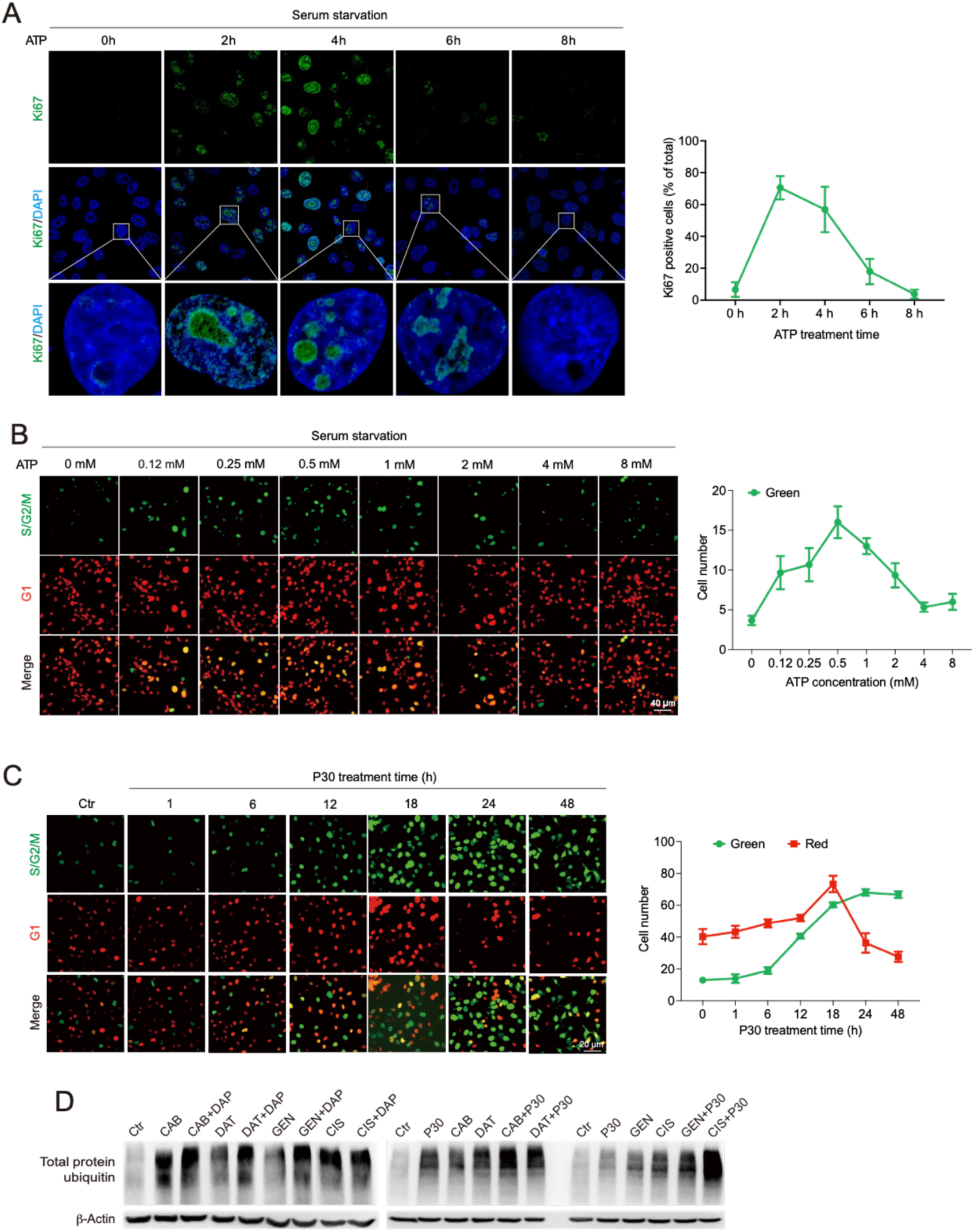
PDK1 inhibition forces ATP production that stimulated cell proliferation. (A) MDA-MB-231 cells in the quiescent state were treated with ATP for indicated time, and the expression of Ki67 was detected by immunofluorescence. (B) MDA-MB-231-CG cells in the quiescent state were treated with increasing dose of ATP, and cell phases were recorded and quantified. (C) MDA-MB-231-CG cells were treated with P30, and cell cycle were recorded and quantified. (D) MDA-MB-231 cells were treated with indicated drugs or combined with PDK1-Is, and protein damage was detected by Western blot. All values are presented as mean value (at least three replications) ± SD, and p value was calculated by comparison with Ctr group or indicated separately.

**Supplementary Fig 10.**
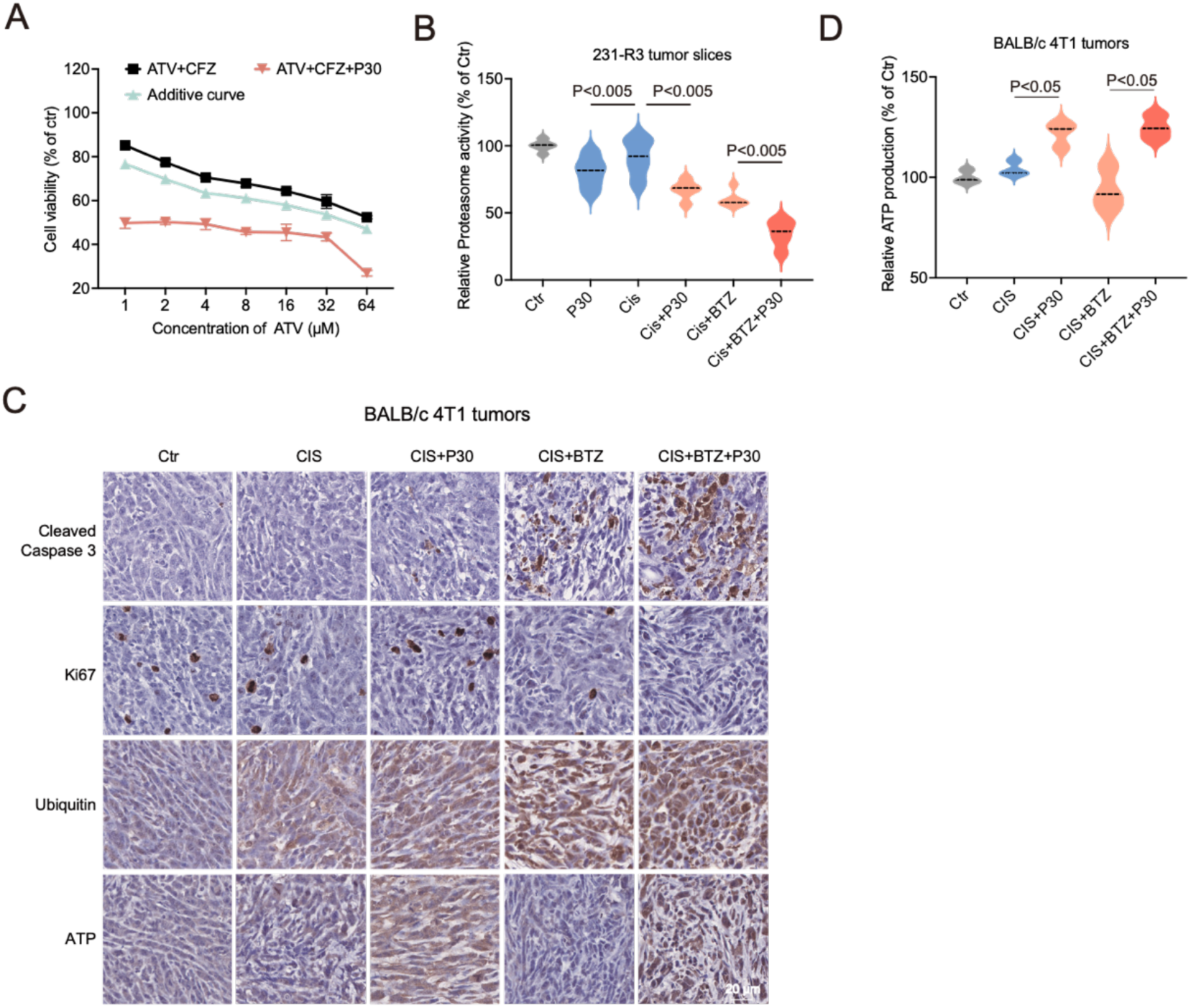
Forced ATP production with proteasome inhibition to overcome MDR. (A) Response of 231-R3 cells to *ATV* combined with non-toxic concentration of proteasome-I or *ATV* combined with non-toxic concentration of proteasome-I and PDK1-I. Cell viability was detected by alamar blue assay. Additive curves were calculated by Additive = E1+E2-E1*E2, where E1 is inhibition effect from *ATV*+*CFZ* and E2 is inhibition effect from P30 treatment. (B) 231-R3 xenograft tumors were cultured in 3D-TSCs and treated with *CIS* or combined with P30 and *BTZ* treatment, each group is quintuplicate, and proteasome activity was measured by fluorescent AMC tagged peptide substrate. (C, D) BALB/c mice bearing 4T1 allograft tumors were treated with vehicle, *CIS* (3 mg/kg), *BTZ* (1 mg/kg), or P30 (1 mg/kg) every 3 days for 3 times, and ATP, cleaved caspase 3, Ki67, and ubiquitin were detected by immunohistochemistry (C). ATP content is also detected by ATP assay kit (D). All values are presented as mean value (at least three replications) ± SD, and p value was calculated by comparison with Ctr group or indicated separately.

**Supplementary Fig 11.**
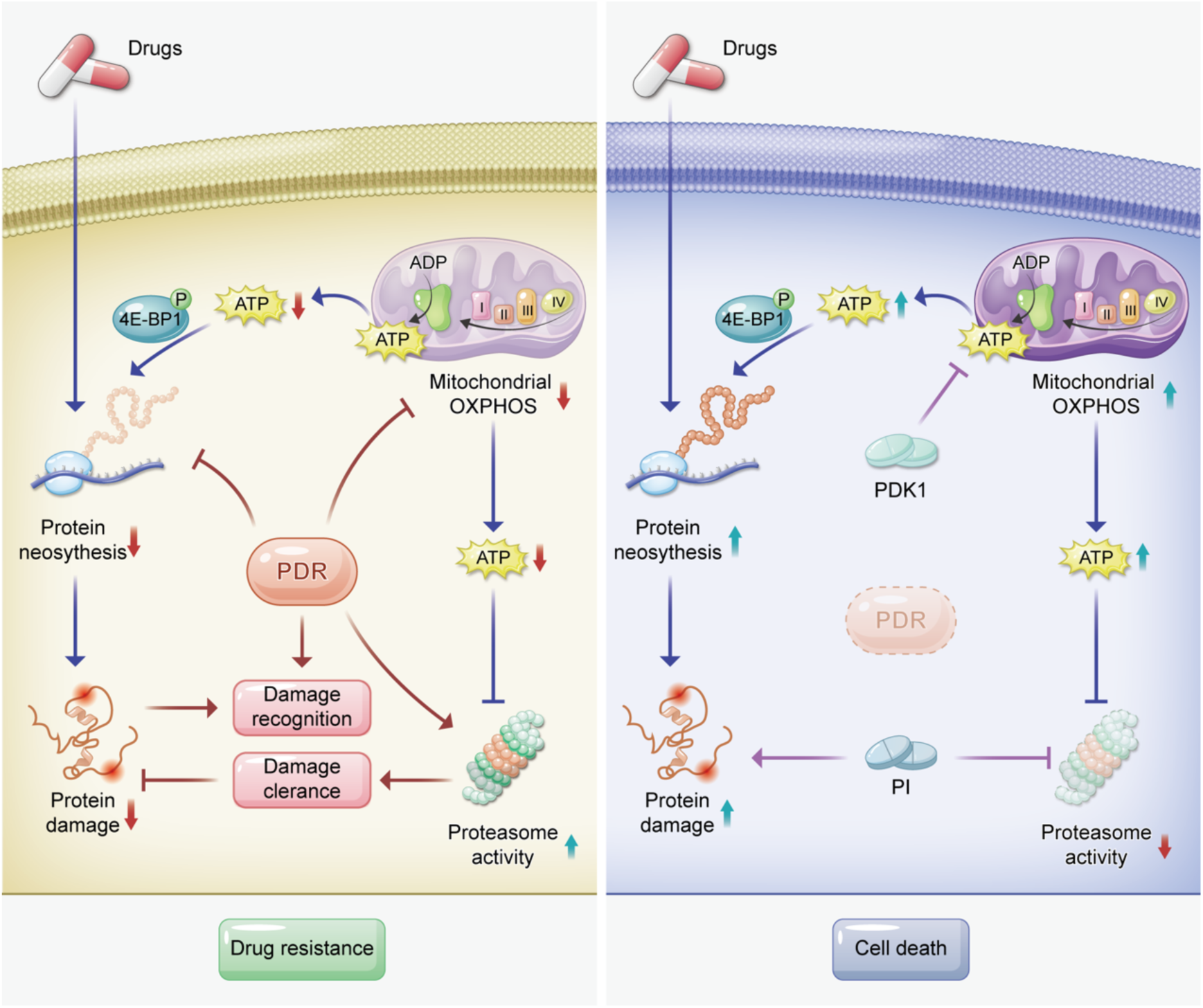
A summary illustrating that majority of anticancer drugs kill cells by damaging newly synthesized proteins and cancer cells initiate a quick response to minimize the cytotoxicity. This study demonstrate that (1) These drugs bind to neosynthesized proteins in a nonspecific fashion and interfere with the proper folding of the bound proteins; (2) The impaired function of damaged proteins should serve as a cause for cytotoxicity of the anticancer drugs; (3) Meanwhile the protein damage also triggers PDR, which includes protein damage recognition by the ubiquitin proteasome system, to evade the lethality which serves as the key mechanism leading to MDR.

