## Supplementary Table 1 for "A critical role of protein damage response in mediating cancer drug resistance"

| Drug Name | Abbreviation | Brand | Cat.# | MDA-MB-231 1h treatment | MDA-MB-231 6h treatment | A549 6h treatment |
|---|---|---|---|---|---|---|
| 10-Deacetylbaccatin | DAB | Selleckchem | S2409 | BD | BD | ND |
| Abiraterone | ABR | Selleckchem | S2246 | BD | BD | BD |
| Abitrexate | ABI | Selleckchem | S1210 | DD | BD | PD |
| Afatinib | AFA | Selleckchem | S1011 | ND | BD | BD |
| Altretamine | ALT | Selleckchem | S1278 | BD | BD | BD |
| Amuvatinib | AMU | Selleckchem | S1244 | BD | BD | BD |
| Anastrozole | ANA | Selleckchem | S1188 | BD | BD | BD |
| Ancitabine | ANC | Sigma | A8598 | BD | BD | BD |
| Artemether | ART | Selleckchem | S2264 | BD | BD | BD |
| Aspirin | ASA | Selleckchem | S3017 | BD | BD | BD |
| Atazanavir | ATV | Selleckchem | S1457 | PD | PD | PD |
| Axitinib | AXI | Selleckchem | S1005 | PD | PD | BD |
| Azacitidine | AZA | Selleckchem | S1782 | BD | PD | BD |
| Azithromycin | AZM | Selleckchem | S1835 | DD | BD | BD |
| Bendamustine | BEN | Selleckchem | S1212 | BD | BD | BD |
| Bepotastine | BEP | Selleckchem | S3037 | PD | PD | BD |
| Bindarit | BIN | Selleckchem | S3032 | PD | BD | PD |
| Bleomycin | BLE | Selleckchem | S1214 | BD | BD | BD |
| Bortezomib | BTZ | BOC Sciences | 179324-69 | PD | BD | BD |
| Bosutinib | BOS | Selleckchem | S1014 | BD | BD | PD |
| Busulfan | BUS | Selleckchem | S1692 | ND | PD | PD |
| Cantharidin | CAN | Sigma | C7632 | ND | BD | ND |
| Carboplatin | CAB | Selleckchem | S1156 | PD | BD | PD |
| Carmofur | CAR | Selleckchem | S1289 | BD | ND | PD |
| Cephalomannine | CEP | Selleckchem | S2408 | PD | ND | BD |
| Chlorambucil | CHL | Selleckchem | S4288 | BD | BD | BD |
| Cisplatin | CIS | Sigma | P4394 | PD | BD | PD |
| Clofarabine | CLO | Selleckchem | S1218 | BD | DD | BD |
| Crizotinib | CRI | Selleckchem | S1068 | ND | DD | PD |
| Cyclophosphamide | CYC | Selleckchem | S2057 | ND | BD | BD |
| Cytarabine | CYT | Selleckchem | S1648 | BD | BD | PD |
| Dacarbazine | DAC | Selleckchem | S1221 | DD | BD | PD |
| Danusertib | DAN | Selleckchem | S1107 | ND | PD | ND |
| DAPT | DAT | Selleckchem | S2215 | PD | PD | DD |
| Dasatinib | DAS | J&K | 923898 | PD | BD | DD |
| Dovitinib | DOV | Selleckchem | S1018 | ND | BD | |
| Eltrombopag | ELT | Selleckchem | S2229 | PD | BD | PD |
| Epirubicin | EPR | J&K | 194237 | BD | BD | BD |
| Erlotinib | ELB | Selleckchem | S1023 | ND | BD | DD |
| Esomeprazole | ESO | Selleckchem | S2233 | PD | PD | BD |
| Estramustine | EMP | Sigma | E0407 | DD | ND | ND |
| Etoposide | ETP | Selleckchem | S1225 | BD | BD | |
| Everolimus | EVE | Selleckchem | S1120 | PD | BD | BD |
| Floxuridine | FLO | Selleckchem | S1299 | BD | BD | BD |
| Fludarabine | FLU | Selleckchem | S1491 | ND | BD | BD |
| Formestane | FMT | Selleckchem | S1300 | PD | PD | BD |
| Fulvestrant | FVT | Selleckchem | S1191 | PD | BD | PD |
| Gefitinib | GEF | Selleckchem | S1025 | PD | DD | PD |
| Gemcitabine | GEM | Selleckchem | S1149 | PD | BD | BD |
| Geniposidic acid | GEA | Selleckchem | S2413 | BD | ND | PD |
| Genistein | GEN | Selleckchem | S1342 | ND | PD | BD |
| Histamine | HIM | Selleckchem | S4118 | PD | PD | PD |
| Ifosfamide | IFO | Selleckchem | S1302 | PD | PD | BD |
| Imatinib | IMA | Selleckchem | S1026 | PD | BD | PD |
| Irinotecan | IRI | Selleckchem | S1198 | BD | BD | BD |
| Itraconazole | ITR | Selleckchem | S2476 | ND | BD | PD |
| Lamotrigine | LTG | Selleckchem | S3024 | PD | BD | PD |
| Lapatinib | LAP | Selleckchem | S1028 | PD | PD | BD |
| Lenalidomide | LEN | Selleckchem | S1029 | ND | BD | PD |
| Letrozole | LET | Selleckchem | S1235 | BD | BD | BD |
| Lomustine | LOM | Selleckchem | S1840 | PD | BD | BD |
| Masitinib | MST | Selleckchem | S1064 | ND | BD | ND |
| Mdv3100 | MDV | Selleckchem | S1250 | DD | DD | BD |

| | | | | | | |
|---|---|---|---|---|---|---|
| Mechlorethamine | MEC | Selleckchem | S4252 | DD | BD | BD |
| Melengestrol | MEL | Selleckchem | E0742 | ND | PD | BD |
| Mercaptopurine | MER | Selleckchem | S1305 | BD | PD | BD |
| Mesna | MES | Selleckchem | S1735 | BD | PD | |
| Methacycline | MCC | Selleckchem | S2527 | BD | BD | BD |
| Methazolastone | MZT | Selleckchem | S1237 | PD | BD | BD |
| Miltefosine | MTF | Selleckchem | S3056 | PD | BD | |
| Mitotane | MTT | Selleckchem | S1732 | PD | PD | ND |
| Mitoxantrone | MXR | Selleckchem | S1889 | DD | DD | DD |
| Moroxydine | MOR | Selleckchem | S2486 | PD | BD | ND |
| Naloxone | NAL | Selleckchem | S3066 | PD | BD | PD |
| Nelarabine | NEL | Selleckchem | S1213 | PD | BD | PD |
| Neratinib | NER | Selleckchem | S2150 | PD | BD | BD |
| Paclitaxel | PTX | Selleckchem | S1150 | BD | BD | BD |
| Paeoniflorin | PAE | Selleckchem | S2410 | BD | BD | BD |
| Pazopanib | PAZ | Selleckchem | S1035 | PD | BD | DD |
| Phenylbutazone | PHB | Selleckchem | S1654 | BD | BD | BD |
| Phenylbutyric | PHE | Selleckchem | S4288 | BD | BD | BD |
| Pioglitazone | PIO | Selleckchem | S2590 | BD | BD | BD |
| Pomalidomide | POM | Selleckchem | S1567 | PD | BD | BD |
| Ponatinib | PON | Selleckchem | S1490 | BD | BD | |
| Proadifen | PRO | Sigma | P1061 | BD | BD | BD |
| Procarbazine | PCB | Selleckchem | S1995 | BD | BD | PD |
| Regorafenib | REG | Selleckchem | S1178 | PD | BD | PD |
| Rosiglitazone | RSZ | Selleckchem | S2556 | PD | BD | PD |
| Ruxolitinib | RUX | Selleckchem | S1378 | ND | DD | ND |
| Sorafenib | SOR | Selleckchem | S1040 | BD | DD | BD |
| Streptozotocin | STZ | Selleckchem | S1312 | PD | PD | PD |
| Sulindac | SUL | Selleckchem | S2007 | PD | PD | PD |
| Temocapril | TEM | Selleckchem | S2099 | PD | PD | PD |
| Teniposide | TEN | J&K | 563738 | BD | BD | |
| Tofacitinib | TFB | Selleckchem | S5001 | PD | BD | DD |
| Tolbutamide | TLB | Selleckchem | S2443 | PD | PD | PD |
| Tolnaftate | TNF | Selleckchem | S2058 | PD | BD | ND |
| Topotecan | TOP | Selleckchem | S1231 | BD | BD | BD |
| Vatalanib | VAT | Selleckchem | S1101 | ND | BD | BD |
| Vemurafenib | VEM | Selleckchem | S1267 | PD | PD | BD |
| Vincristine | VIN | Selleckchem | S1241 | BD | PD | BD |
| 2-Methoxyestradiol | MTD | Selleckchem | S1233 | | BD | |
| Adrucil | ADR | Selleckchem | S1209 | | BD | |
| Carbazochrome | CZC | Selleckchem | S3000 | | PD | |
| Capecitabine | CAP | Selleckchem | S1215 | | PD | |
| Docetaxel | DCT | Selleckchem | S1148 | | BD | |
| Doxifluridine | DFD | Selleckchem | S2045 | | BD | |
| Doxorubicin | DOX | Selleckchem | S1208 | | BD | |
| Vinorelbine | VRB | Selleckchem | S4269 | | BD | |
| Carmustine | CMS | Sigma | C0400 | | ND | |
| Cladribine | CLA | Selleckchem | S1199 | | DD | |
| Dacomitinib | DMT | Selleckchem | S2727 | | BD | |
| Flutamide | FTM | Selleckchem | S1908 | | ND | |
| Ftorafur | FTO | Selleckchem | S1300 | | DD | |
| Temsirolimus | TLM | Selleckchem | S1044 | | BD | |
| Azaguanine-8 | AGN | Selleckchem | S4194 | | | BD |
| Bergapten | BER | Selleckchem | S4239 | | | BD |
| Cephalomannine | CEP | Selleckchem | S2408 | | | BD |
| Cinacalcet | CIN | Selleckchem | S1260 | | | DD |
| Clorsulon | CSL | Selleckchem | S2613 | | | PD |
| Neratinib | NER | Selleckchem | S2150 | | | BD |
| Nilotinib | NIL | Selleckchem | S1033 | | | BD |
| Nilvadipine | NVP | Selleckchem | S2721 | | | PD |
| Oxaliplatin | OXA | Selleckchem | S1224 | | | BD |
| Pemetrexed | PEM | Selleckchem | S7785 | | | DD |
| Saracatinib | SAR | Selleckchem | S1006 | | | PD |
| Tretinoin | TRE | Selleckchem | S1653 | | | ND |
| Vandetanib | VAT | Selleckchem | S1046 | | | BD |
| Vismodegib | VIS | Selleckchem | S1082 | | | BD |
| Vorinostat | VOR | Selleckchem | S1047 | | | BD |

| | | | | |
|---|---|---|---|---|
| XI-184 | XIA | Selleckchem | S1119 | BD |
| Zoledronic | ZOL | Selleckchem | S1314 | BD |
| Toremifene | TOR | Selleckchem | S1776 | PD |
